# Coral genetic structure in the Western Indian Ocean mirrors ocean circulation and thermal stress

**DOI:** 10.1101/2025.06.16.659828

**Authors:** Annie S. Guillaume, Stéphane Joost, Sarvanen Curpen, Danishta Dumur Neelayya, Luxmibye Harree-Somah, Oocheetsing Sadasing, Luca Saponari, Charlotte Dale, Léo Barret, Nina Andrews, Sanjeev Kumar Leckraz, Ronnie François, Vasisht Seetapah, Vinayaganidhi Munusami, Suraj Bacha Gian, Reshad Jhangeer-Khan, Terence Mahoune, Pramod Kumar Chumun, Manuel Poretti, Véronique Berteaux-Lecellier, Gael Lecellier, Oliver Selmoni

## Abstract

Global warming and rising sea temperatures are pushing many reef-building coral species towards extinction. As thermal tolerance in corals is partially heritable, identifying genes under thermal selection is critical for targeted biodiversity management. However, it remains unclear how large breaks in connectivity (>100 km of open sea) affect the spread of adaptive alleles for different coral species in discontinuous reef networks such as the West Indian Ocean (WIO). To address this, we applied a seascape genomics approach to model (i) population connectivity and (ii) thermal adaptive potentials for two keystone coral species, *Acropora muricata* and *Pocillopora damicornis*, across the WIO. For both species, corals from the Seychelles were predominantly genetically isolated from corals in Rodrigues and Mauritius, putatively an effect of regional oceanographic barriers. Furthermore, sea currents during reproductive periods better predicted genetic connectivity than did Euclidean distances for both species, highlighting that connectivity models can serve as proxies to understand dispersal potential depending on reproductive strategies. Spatial patterns of neutral genetic variation were best explained by sea surface temperature variability and mean degree heating weeks. When used in genotype– environment association (GEA) analyses, we identified hundreds of loci under putative thermal selection from linked to known heat stress responses. In *A. muricata*, five Sacsin genes—co- chaperones of the Hsp70 heat-shock protein involved in thermal stress response—were identified, alongside genes related to immune defence, antioxidant response, signalling, and protein folding. In contrast, only the centromere protein V, involved in mitosis, was enriched in *P. damicornis*. By integrating patterns of gene flow with molecular adaptations to estimate species-specific adaptive potentials, we found that large sea distances and strong oceanographic barriers inhibit the genetic exchange of adapted genotypes across the WIO, providing valuable insights to guide local and regional biodiversity management in this region.

## 1. Introduction

Covering less than 1% of the ocean floor, coral reefs are biodiversity hotspots hosting approximately a third of all marine species (Moberg and Folke, 1999). These dynamic ecosystems provide valuable habitat for a variety of taxa, shaping food chains and biogeochemical cycles while also providing important ecosystem services for humans, including food, medicine, coastal protection, and tourism (Costanza et al., 2014; Hughes et al., 2017a; Moberg and Folke, 1999; Roff et al., 2016). The foundation of these ecosystems are reef-building scleractinian corals, whose calcareous skeletons construct the reef’s physical structure (Hughes et al., 2017a; Moberg and Folke, 1999). Coral reefs are under threat of intensifying anthropogenic-induced stressors, particularly from global warming and elevated sea temperatures (Bozec et al., 2025; Hughes et al., 2020, 2017b; Ortiz et al., 2021; Otto, 2018; Virgen-Urcelay and Donner, 2023). Under such stressful conditions, corals can lose their symbiotic algae in a reversible process known as coral bleaching (Helgoe et al., 2024; Hoegh- Guldberg et al., 2007). As corals rely on their symbionts for metabolic inputs, persistent bleaching can result in widespread coral deaths (Sully et al., 2022). An estimated 14% of the world’s coral cover has already been lost between 2009 and 2018 (Souter et al., 2021), and with bleaching-induced coral mortality on the rise (Bozec et al., 2025; Hughes et al., 2020, 2017b; Virgen-Urcelay and Donner, 2023) there is concern for the impact of climate change on coral reef structures, biodiversity, functioning and productivity globally (Graham et al., 2015; Hughes et al., 2019; Morais et al., 2022; Sully et al., 2022).

Despite increased frequency and intensity of mass bleaching events, corals appear to be adapting to repeated exposure of elevated temperatures (e.g., Bozec et al., 2025; Drury and Lirman, 2021; Louis et al., 2016; Palumbi et al., 2014; Schoepf et al., 2015). For example, increased thermal tolerance of coral communities has been observed around Palau in the Pacific Ocean in recent years following multiple acute bleaching events across the last few decades (Bruno et al., 2001; Lachs et al., 2023). While these trends could be attributed to acclimatisation, shifts in community structure or changes in the holobiont associations (Gouezo et al., 2019), contemporary within-population variation in coral heat tolerance indicates the possibility of an underlying genetic adaptation to elevated temperatures (Lachs et al., 2023; Mumby and Van Woesik, 2014). Indeed, studies using a variety of experimental methods (summarised in Selmoni et al., 2024) found that within-species thermal variation is heritable, with evidence arising from the Pacific Ocean (e.g., Bay and Palumbi, 2014; Selmoni et al., 2021, 2020a), the Great Barrier Reef of eastern Australia (e.g., Cooke et al., 2020; Dixon et al., 2015; Elder et al., 2022; Fuller et al., 2020; Jin et al., 2016; Lundgren et al., 2013; Quigley et al., 2020), the East Indian Ocean along the west Australian coast (e.g., Thomas et al., 2022, 2017), the Persian Gulf in the Middle East (e.g., Howells et al., 2021, 2016; Kirk et al., 2018; Smith et al., 2022), and the Caribbean Sea (e.g., Drury and Lirman, 2021; Dziedzic et al., 2019). As the heritability of thermal variation implies the existence of alleles that can promote tolerance to anticipated increased thermal regimes, identifying candidate genes and populations adapted to thermal stresses can provide valuable information for improving spatial mapping of reef vulnerability to inform reef restoration efforts (Selmoni et al., 2024).

Seascape genomics provides a promising framework for investigating adaptation to environmental stressors in *in-situ* populations located across diverse geographic and environmental ranges (Riginos et al., 2016; Selmoni et al., 2021, 2020a). Within this framework, neutral population structuring and connectivity across large spatial extents can provide key insights for understanding how sea currents might affect the spread of adaptive alleles within and between coral populations (De Mita et al., 2013; Lotterhos and Whitlock, 2015; Matz et al., 2020; Selkoe et al., 2016; Selmoni et al., 2020a; Weersing and Toonen, 2009). Seascape genomics also provides a framework for correlating whole genome variation with environmental factors using genotype–environment associations (GEA). In this way, genes putatively involved in adaptation are identified while simultaneously providing insights into the environmental conditions that might be driving genetic differences (Rellstab et al., 2015). Significant associations from GEA correlations suggest candidate loci under selection, which, if reference genomes are available, can be used to highlight markers near or within a gene of interest to infer molecular functions under selection (Storfer et al., 2018). This is important, as long-term coral cover is expected to benefit from genetic influxes of thermally-adapted coral recruits originating from reefs experiencing temperatures 0.5°C higher than the local average (Matz et al., 2020). To assist the long-term adaptive capacities of coral populations, it is imperative to understand connectivity between reefs and the genes involved in adaptation.

While strides are being made to understand coral thermal adaptations and thermal heritability globally, research gaps remain. First, research is predominantly concentrated around reef systems in the Indo-Pacific region, notably the Great Barrier Reef in Australia (summarised in Selmoni et al., 2024). For instance, early GEA methods successfully detected candidate genes under thermal selection in the Great Barrier Reef (Lundgren et al., 2013). However, large stretches of coral reef regions remain understudied despite coral bleaching responses varying within and among oceans, attributable to differences in regional thermal regimes, evolutionary histories, and past disturbances (Shlesinger and van Woesik, 2023). Notably, coral reefs in the West Indian Ocean (WIO) have been largely overlooked (Carr et al., 2025), despite hosting approximately 5% of global coral reefs affected by recurrent short- and long-term bleaching events (Shlesinger and van Woesik, 2023; van Woesik and Kratochwill, 2022). With most WIO ecosystems labelled as vulnerable (e.g., Seychelles) to critically endangered (e.g., Mauritius Islands) due to human activities and global warming (Obura et al., 2022), more research is necessary in this region for informed biodiversity management. Second, research typically focuses either on contiguous reef systems or at local isolated reefs (Selmoni et al., 2024).

Though marine environments are highly connected in general, where local stressors can have regional-level effects on community structures (Green et al., 1987), it remains unclear of how breaks in connectivity (>100 km of open sea) affect the spread of adaptive alleles between reefs of a network. Last, research into coral thermal adaptation is typically conducted on a single study species at a time (Selmoni et al., 2024; but see Selmoni et al., 2021). Diversity of morphologies and habitat preferences results in differential responses of corals to selection pressures and stressful events (Guest et al., 2012; Johnston et al., 2021; Yuen et al., 2023). For example, corals with branched morphologies are generally more sensitive to heat stress than those with massive morphologies (Loya et al., 2001). As evolutionary trajectories are species-specific, it seems that patterns of genomic diversity observed in one species cannot necessarily be extrapolated to others, yet conservation strategies must consider the needs of multiple species simultaneously (Voolstra et al., 2023).

Despite a growing body of research investigating coral connectivity between islands of the WIO (e.g., Carr et al., 2025; Gélin et al., 2018; Kusumoto et al., 2020; Oury et al., 2024; Vogt-Vincent et al., 2024), studies have yet to incorporate genetic data for investigating region-wide connectivity between Seychelles and the Mascarene Islands, with little investigation into the adaptive potentials of corals across the WIO region. Here, we aim to address these research gaps to better understand coral adaptive potential across large, non-contiguous reef networks. Specifically, we investigate (i) the genetic connectivity of two keystone coral species with contrasting reproductive strategies across the WIO, and (ii) their molecular adaptations to thermal conditions. To this end, we applied a seascape genomics framework to detect coral reefs that putatively host thermal stress-adapted coral genotypes, to ultimately provide information that will support local stakeholders in developing regional biodiversity management plans in the WIO (Donovan et al., 2021; Goetze et al., 2021; Pittman et al., 2021; Stefanoudis et al., 2023).

## 2. Materials and Methods

We focus on two common WIO reef-building coral species, *Acropora muricata* and *Pocillopora damicornis*, sampled at reefs around the Seychelles in the northern WIO and Mauritius and Rodrigues of the Mascarene Islands group in the southern WIO (**Fig. 1a**). We first characterised the seascape conditions around each island using open-access satellite and geomorphic data, from which 15 reefs with contrasting environmental conditions were selected for coral sampling and genotyping. Our framework follows a seascape genomics approach (Selmoni et al., 2020a) to (i) map the connectivity of populations across the WIO region and (ii) identify putatively adaptive coral genotypes using genotype–environment association (GEA) analyses (**Suppl. Fig. 1**). By coupling these analyses, we then infer the spatial distribution of WIO reefs that potentially carry genotypes underpinning local adaptation to thermal stress for informed reef management planning. All analyses were performed in the R environment (R Core Team, 2024).

**Figure 1.**
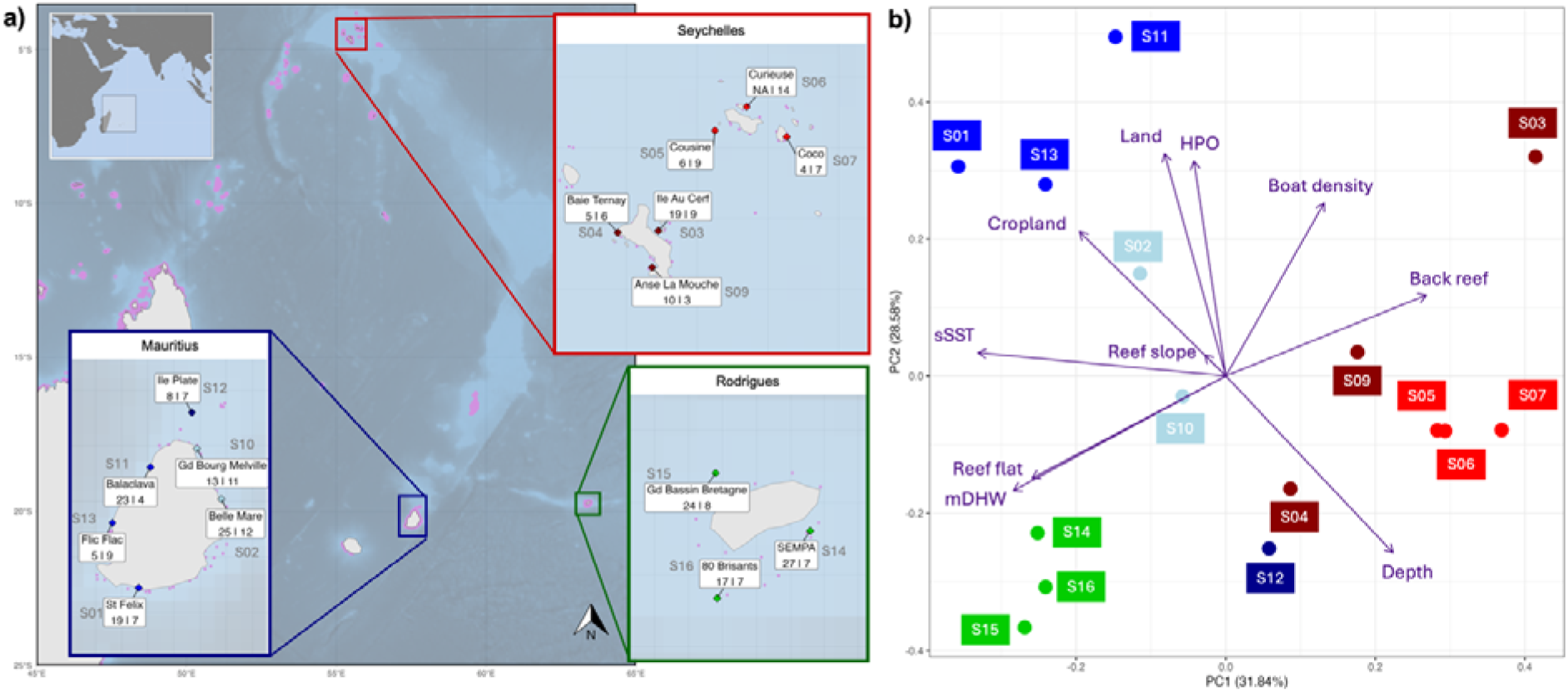
**a)** Map of the study area in the West Indian Ocean (WIO; map insert, top right) indicating the 15 sampling sites distributed around the Seychelles (Mahé: dark red, Praslin: red), and two of the Mascarene Islands of Mauritius (blue) and Rodrigues (green). Names and site identifiers (S#) used in this study are provided for each reef. The number of genotyped individuals retained after quality filtering are indicated at each sampling site per species, with values first for *Acropora muricata* then *Pocillopora damicornis*. **b)** Environmental characterisation of the study reefs using 10 uncorrelated environmental variables (**Table 1**), where abbreviations are as follows: sSST = standard deviation of sea surface temperature; mDHW = mean degree heating week; HPO = human population density.

**Table 1.** Summary of the uncorrelated environmental variables used to characterise the seascape of 15 reefs sampled for this research.

| Variable | Acronym | Source | Spatial resolution | Temporal resolution |
| --- | --- | --- | --- | --- |
| Depth | DEP | RECIFS | 5x5 km | n/a |
| Degree Heating Week (mean) | mDHW | RECIFS | 5x5 km | Monthly max from 1985–2021 |
| Sea Surface Temperature (standard deviation) | sSST | RECIFS | 5x5 km | Monthly max from 1985–2021 |
| Density of cropland | CROP | RECIFS | 5x5 km | Measured in 2015–2019 |
| Density of land | LAND | RECIFS | 5x5 km | n/a |
| Human population along coastline | HPOP | RECIFS | 5x5 km | Measured in 2000, 2005, 2010, 2015, 2020 |
| Boat detection | BOAT | RECIFS | 5x5 km | Measured in 2017–2021 |
| Fraction of reef slope in 250m around reef | ReefSlope | ACA | 10x10 m | n/a |
| Fraction of reef flat in 250m around reef | ReefFlat | ACA | 10x10 m | n/a |
| Fraction of back reef in 250m around reef | BackReef | ACA | 10x10 m | n/a |

### 2.1 Study species

We selected *A. muricata* and *P. damicornis* for the present study as two common reef-building corals of the WIO that represent keystone species with high ecological relevance (Bridge et al., 2023; Cunning et al., 2018). Both species show evidence of population recovery after severe bleaching events (Bruno et al., 2001; Lachs et al., 2023), which could be attributed to thermal tolerance (Ainsworth et al., 2016; Carr et al., 2025; Gouezo et al., 2019; Humanes et al., 2022), such that they are interesting subjects for investigating adaptation to thermal stressors.

Furthermore, these species have distinct life history strategies that warrant independent investigations of thermal tolerance and connectivity. *A. muricata* is a predominately broadcast spawning coral, releasing gametes into the water column in synchronised events at the beginning of the Austral summer (October to December) when sea temperatures increase (Wijayanti et al., 2019). *P. damicornis* has mixed reproduction, with sexual reproduction via year-round broadcast spawning (Gélin et al., 2018, 2017; Schmidt-Roach et al., 2012; Smith et al., 2019; Ward, 1992).

### 2.2 Sampling strategy

Fifteen reefs were selected for sampling across three regions of the WIO: six sites around Mauritius, six sites around the Seychelles (three each at Mahé and Praslin) and three sites around Rodrigues (**Fig. 1a**; **Suppl. Table 1**). Reefs were selected following the methods of Selmoni et al. (2020b) to maximise environmental contrast for improved power in seascape genomic analyses, while simultaneously considering logistical constraints in collaboration with local stakeholders, including site accessibility and weather conditions during sampling. Full details are provided in the Supplementary Material (**Suppl. Methods: Selection of sample sites**).

The field sampling campaign took place with the assistance of scientists and technical staff of the Mauritius Oceanography Institute (MOI) in January and February 2022 for Mauritius and Seychelles, and in May 2022 for Rodrigues. This sampling campaign was performed within the framework of the Coral Reef Restoration Project of the United Nations Development Programme Mauritius (PIMS 5736, sampling permit in Rodrigues SPA 30) and the Government of Seychelles (sampling permit A0157). Colonies identified as *A. muricata* and *P. damicornis* were sampled for genotyping within a radius of 250m from the coordinates of the sample site at depths between 2-10m, with up to 30 colonies sampled per site per species. Each sample consisted of a 2cm branch collected with pliers and bagged underwater, before being transferred to 80% ethanol and stored at -20°C. A total of 345 *A. muricata* and 403 *P. damicornis* colonies were sampled for DNA extraction and single nucleotide polymorphism (SNP) genotyping (**Suppl. Table 1**).

### 2.3 Environmental characterisation

The seascapes at the 15 study reefs were characterised using 17 environmental and geomorphic variables that might be exerting selection pressures on corals. These variables were accessed from two publicly available online archives. Environmental variables pertaining to climatic conditions and proximity to anthropogenic activities covering the whole study extent were extracted from the Reef Environment Centralized Information System (RECIFS; Selmoni et al., 2023). These variables were accessed at 5 km^2^ spatial resolutions and described as month- by-month environmental variation covering a temporal range of a minimum of 20 years up to December 2021 (i.e., when sampling took place; full details on variables are summarised in **Suppl. Table S2**). Temporal mean and standard deviations were then calculated for degree heating week (DHW; Skirving et al., 2020), sea surface temperature (SST; Skirving et al., 2020), chlorophyll (CHL) and suspended particulate matter (SPM), where CHL and SPM were obtained from the ‘OCEANCOLOUR_GLO_BGC_L4_MY_009_104’ dataset (accessed on 03-04-2014; E.U. Copernicus Marine Service Information (CMEMS), 2024). In addition to RECIFS environmental variables, we retrieved a geomorphic characterisation of the WIO reefs from the Allen Coral Atlas (ACA; Allen Coral Atlas, 2022). Available at 10m resolution, this variable was processed to calculate the proportion of reef slope, reef flat, back reef and reef plateau at each sampling site (250m radius around the sample site coordinates). A Spearman’s pairwise non-parametric rank correlation threshold of ρ<|0.8| was applied to retain uncorrelated variables for downstream analyses (Table 1), where a principal component analysis (PCA) on scaled and centred environmental variable values was used to characterise the study sites (**Fig. 1b**).

### 2.4 DNA extraction and SNP genotyping

DNA extraction and SNP genotyping followed methods of Selmoni et al. (2021). Briefly, DNA was extracted using a DNeasy Blood and Tissue 96 kit (Qiagen) following the manufacturer’s instructions. DNA samples were sent to Diversity Arrays Technology (Canberra, Australia) for quality check screening and genotype-by-sequencing using the DArT-sequencing method (DArT-seq), using restriction enzymes *Pst*I and *Hpa*II (Gawroński et al., 2016; Kilian et al., 2012; Selmoni et al., 2021). Sequencing read processing was performed by Diversity Arrays Technology using proprietary DArT analytical pipelines (Sansaloni et al., 2011). To minimise technical bias and potential batch effects, targeted *A*. *muricata* and *P. damicornis* samples were kept separated and randomly distributed across the respective batches (e.g., 96-well plates, sequencing lanes) throughout the workflow (DNA purification, library preparation and sequencing). Raw sequence data have been deposited in the NCBI BioProject database under accession number PRJNA1277000.

### 2.5 SNP filtering

The raw DArT-seq loci were aligned to each species’ reference genomes using BLAST to retain only the SNPs associated with the coral hosts (filtering thresholds: >70% percentage identity, >80% overlap identity, and >50 bitscore). The reference genome for *A. muricata* was the chromosome-level assembly of *Acropora millepora* (v2.1; GCF_013753865.1; Fuller et al., 2020), and the reference genome for *P. damicornis* was the scaffold-level assembly of *P. damicornis* (v1; GCF_003704095.1; Cunning et al., 2018).

SNP filtering was performed for each species separately. First, we filtered out individuals sharing highly similar genotypes (Pearson pairwise correlations >0.95), retaining one individual in cases where genetic similarity was found. We then removed SNPs and individuals with high missing rates (>80%) before applying a filter to exclude rare alleles (minor allele frequency; MAF <5%). As both species potentially include cryptic species (Gélin et al., 2018, 2017; Grupstra et al., 2024; Johnston et al., 2017; Oury et al., 2024; Riginos et al., 2024), we used a principal coordinate analysis (PCoA) of the genotype matrix to cluster individuals based on genomic distance, before removing clusters with an F_ST_ difference >0.3 with the other clusters. These individuals will hereafter be referred to as ‘cryptic’ individuals. We genetically confirmed the species identity of retained *Pocillopora* samples as *P. damicornis* using DArT sequences within the ITS-2 and 18S ribosomal DNA regions (**Suppl. Table S3**). We repeated the missingness and MAF filtering steps on the retained individuals. Last, we assessed linkage disequilibrium (LD) amongst SNPs using functions from the *dartR* R package (Mijangos et al., 2022) to create an LD map of the genome, with a maximum pairwise limit of 15 kbp (the distance of estimated LD decay to r^2^<0.05, based on whole genome sequence data of *A. millipora*; Fuller et al., 2020) before pruning SNPs with an LD >0.2. As we found minimal effect of LD pruning on population structure (assessed using a PCoA), nor evidence of SNPs clumping in a PCA (assessed using *PCAdapt*; Luu et al., 2017; Privé et al., 2020), we did not prune for LD and retained all loci for downstream analyses.

### 2.6 Neutral genetic structure

We assessed the neutral genetic structure of each species using two complementary methods: i) Principal Coordinate Analysis (PCoA) and ii) Sparse Nonnegative Matrix Factorization (sNMF).

We first produced a neutral genomic dataset, where we identified and removed outlier loci using a genome scan of the filtered genotype matrix via the *pcadapt* R package (v4.3.5; Luu et al., 2017; Privé et al., 2020). In short, loci were attributed p-values based on their Mahalanobis distances along the retained axes of a PCA of the genotype matrix (2 PCs for *A. muricata* and 3 PCs for *P. damicornis*). We calculated the false discovery rate (FDR; Storey and Tibshirani, 2003) to correct p-values for multiple testing using the *qvalue* R package (2.36.0; Storey et al 2024), with a q-value <0.05 threshold to identify and remove outlier loci for the neutral genotype matrix.

We assessed contemporary population structure using a Gower PCoA clustering mixture model through the *dartR* R package (v2.9.7; Gruber et al., 2018; Mijangos et al., 2022) based on a distance matrix from allele frequency differences amongst individuals. We retained 2 PCo axes for *A. muricata* and 3 PCo axes for *P. damicornis* before allocating individuals to population groups using a hierarchical cluster analysis. We assessed ancestral population structure using sNMF algorithms with the *LEA* R package (v3.16.0; Frichot and François, 2015). Using a cross- entropy validation for K=1–10 over five repetitions with an alpha=100, we identified the number of ancestral populations to retain, using the K values resulting in the lowest average cross entropy to compute the sNMF. Membership assignment of each individual to the K ancestral populations was represented as proportions in a barplot. We calculated genetic structure between the sample sites using *F-*statistics (Wright, 1969, 1978) through the *dartR* R package (Gruber et al., 2018; Mijangos et al., 2022), obtaining observed (H_O_) and expected (H_E_) heterozygosity levels to calculate the inbreeding coefficient (F_IS_), alongside pairwise F_ST_ between sample sites.

### 2.7 Connectivity analysis

We ran a connectivity analysis to investigate how physical distances between reefs correspond to genetic separation between colonies, calculated using the pairwise F_ST_ index (Section 2.6).

Physical distance between reefs were derived from sea current data, following methods of Selmoni et al. (2020a). Briefly, we retrieved maps describing monthly average direction and strength of surface sea currents throughout the WIO. These maps are available at a spatial resolution of 0.083° across 30 years as satellite-derived reconstructions of sea currents (from the ‘GLOBAL_REANALYSIS_PHY_001_030_104’ dataset, accessed on 03-04-2014; E.U. Copernicus Marine Service Information (CMEMS), 2024), which are publicly available via RECIFS (Selmoni et al., 2023). For each pixel, we calculated the cumulative speed towards each of the eight neighbouring pixels and divided this by the total speed to obtain a probability of transition in each direction (the conductance). We then calculated the dispersal costs as the inverse of the square conductance to obtain transition matrices using the *gdistance* R package (v1.6.4, van Etten, 2017). Finally, we calculated the shortest sea distance (least-cost path) between pairs of sites in both directions from the transition matrix to obtain an asymmetrical square matrix of shortest sea distance.

To determine the best predictor of genetic distance between reefs, we built a set of linear connectivity models to estimate F_ST_ as a function of the shortest in-water sea distance between sample sites. We built one model per month, and one based on annual average sea currents (13 models total). We also investigated the predictive performance of Euclidean geographic distances as an independent comparison model calculated from cartesian coordinates of sampled reefs.

Models were built using a log-log relationship between F_ST_ and physical distance (i.e., sea distance and geographic distance). Their predictive powers were assessed using a leave-one- reef-out jackknife resampling approach, where model performance was assessed using coefficients of determination (R^2^). Predictions of genetic connectivity among sites were obtained by translating the unit of sea distance into a unit of genetic separation (F_ST_) through a linear model, using the connectivity predictor that best explained F_ST_ values.

### 2.8 Genotype–environment associations (GEA)

We performed GEAs using multivariate redundancy analyses (RDA) at the population-level following methods of Capblancq and Forester (2021). The genotype response matrices were coded as allele frequencies, where missing data was imputed from the median of the locus- specific allele frequency across all populations (missingness per site for *A. muricata*: S04 with 0.05%, S05 with 0.04%, S07 with 0.14%; and for *P. damicornis*: S04 with 0.01%, S09 with 0.06%). The environmental variables used in the explanatory matrix were selected from the 10 uncorrelated variables (Table 1) using an autonomised forward and backward stepwise model selection procedure with 10,000 permutations through the *ordistep* function of the *vegan* package. We first ran a preliminary RDA to identify environmental variables explaining neutral genomic variation. This procedure selected the standard deviation of monthly sea surface temperature (sSST) and average monthly maximal DHW (mDHW) as the variables that best explained neutral genetic variation independently for both species. To detect putative genomic regions under selection, we performed a multivariate GEA using a partial RDA on the entire genotype matrix and included reef-level population structure as a conditional variable to control for neutral genetic variation. Controlling for neutral genetic variation was done to reduce false- positive detections at the expense of losing power to detect true outlier loci along neutral gradients (Excoffier et al., 2009; Lotterhos, 2023). The values representing population structure were obtained from the axes of a PCA of centred and scaled neutral allele frequencies using the *rda* function of the *vegan* R package, retaining the first *K* PCs that explained more than the mean explained variance.

Outlier loci were identified from RDA loadings (Capblancq et al., 2018). After retaining all constrained RDA axes (*K*=2 for both species), we evaluated the significance of each SNP based on the extremeness of its Mahalanobis distance value compared to the distribution of the other SNPs in the RDA space. The Mahalanobis distances were computed using the *dist_ogk* function of the *bigutilsr* R package (v.0.3.4; Privé, 2021), corrected for genomic inflation factor (François et al., 2016) and distributed along a chi-squared distribution with *K* degrees of freedom to assign a p-value to each SNP (Luu et al., 2017). We applied an FDR threshold of *q*-value<0.05 to identify outlier loci.

### 2.9 Gene Ontology (GO) enrichment analysis

Gene Ontology (GO) enrichment analyses were used to assess the putative molecular function(s) of significant outlier SNPs (i.e., those with *q*-value <0.05) detected with the two heat-related variables (sSST and mDHW) for both species, following Selmoni et al. (2021). To facilitate comparability of annotation between species, we re-annotated genes in both reference genomes. For this, we retrieved transcript sequences of each gene, then ran a similarity search (blastx) against a manually curated database of protein sequences and annotations (Uniprot/swissprot, metazoan entries as accessed on November 2022; Boeckmann et al., 2003). Protein annotations were assigned to the coral genes when the E-score of the similarity search was <10^-6^. We then identified genes located within ±10 kbp of significant SNPs from the GEA analysis (hereafter “significant genes”; Selmoni et al., 2021). We evaluated gene-set enrichment of significant genes using the *SetRank* R package (version 1.0; Simillion et al., 2017), where genes are ranked based on ascending *q*-values from the GEA from which overrepresented GO terms associated with significant genes are identified. GO terms were deemed significant when they had *SetRank p-*values<0.01 and adjusted *p*-values<0.05 (i.e., corrected for multiple testing and overlap between gene-set categories).

### 2.10 Adaptive Potential Index (API)

The potential of corals to adapt to temperature increases depends on (i) the thermal history at the reef (and therefore local adaptation of populations) and (ii) the arrival of propagules from neighbouring heat-adapted reefs (Matz et al., 2020; Selmoni et al., 2020a). Building on previous adaptation indices, including the Population Adaptive Index (Bonin et al., 2007) and the Landscape Adaptive Index (Zhang et al., 2019), we calculated a spatially explicit ‘Adaptive Potential Index’ (API) by combining results from the GEA (Section 2.7) and the connectivity analyses (Section 2.9). To summarise potential for local adaptation, first we calculated an ‘Adaptive Score’ to link putatively adapted alleles with the thermal history of reefs across the WIO (Capblancq et al., 2020). For this, we ran an RDA associating the outlier loci from the GEA (Section 2.7) with the thermal explanatory variables (i.e., sSST and mDWH). The environmental scores from these RDA axes were extracted and normalised to a scale of 0-1 to obtain an Adaptive Score (Steane et al., 2014). The Adaptive Score was then projected across the WIO reefs using thermal history data for each reef of the region. Second, we calculated the ‘Inbound Connectivity Index’ (ICI) and ‘Outbound Connectivity Index’ (OCI) for each reef, which represent the total neighbouring reef areas located downstream or upstream of a target cell, respectively, limited to a sea distance equivalent to the species’ F_ST_ ≤0.1 (Section 2.9). Finally, we calculated the Adaptive Potential Index for each reef cell as the total area of inbound connected reefs with an Adaptive Score >0.8.

## 3. Results

### 3.1 Site characteristics and genomic filtering

Across the 15 reefs selected for coral sampling, six sites were sampled in the Seychelles (three each at Mahé and Praslin), six in Mauritius and three in Rodrigues (**Fig. 1a**). These reefs represent contrasting environmental conditions (**Fig. 1b**), as characterised by 10 uncorrelated environmental variables (Table 1; **Suppl. Fig. S2**). The northern-most sampled reefs were those from the Seychelles (represented in red), comprising predominantly of back reefs located near deep water with less thermal stress (i.e., low sSST and low mDHW), generally further away from human activity (i.e.., cropland, human population and boat activity). Most Mauritian reefs (represented in blue) are characterised by reef flats near land and human activity (i.e., cropland and boats) that experience more variable thermal conditions. The exception is the more isolated northern-most Mauritian site on Ile Plate (S12 in dark blue), which is associated with deeper waters and more stable temperatures. Finally, the three reefs at Rodrigues (represented in green) are isolated reef flats located near deeper water that experience hotter (higher mDHW) and more variable SST conditions (sSST).

Between 8–29 individuals per species were sampled for genotyping at each site (**Fig. 1a**; **Suppl. Table S1**). The DArT-seq analytical pipeline resulted in 73,253 bi-allelic SNPs genotyped for 345 *A. muricata* individuals and 65,708 SNPs for 403 *P. damicornis* individuals (values of SNPs and individuals retained at each step are summarised in **Suppl. Table S4**). We removed 48 *A. muricata* and 261 *P. damicornis* individuals identified as clones, followed by 49 *A. muricata* and 8 *P. damicornis* individuals with too much missing data and low MAF, and a further 43 *A. muricata* and 14 *P. damicornis* individuals identified as cryptic individuals (i.e., F_ST_>0.3 between PCoA clusters). We used pairwise F_ST_ and PCoA biplots to confirm that these cryptic individuals were removed from the dataset (**Suppl. Fig. S3**). After filtering for rare variants, missing values, clones and cryptic individuals, our final genotype matrix comprised of 205 *A. muricata* individuals with 5,757 SNPs and 120 *P. damicornis* individuals with 12,953 SNPs. This resulted in 4–7 *A. muricata* individuals retained from 14 sites (no individuals were left at Curieuse Reef (S06) in Seychelles), and 3–14 *P. damicornis* individuals retained across all 15 sites (**Suppl. Table S1**).

### 3.2 Neutral genetic structure

Outlier loci were identified and removed using a genome scan via the *pcadapt* R package to produce a neutral genotype matrix for assessing population structure across the WIO. For the genome scan, we retained *K*=2 PCs for *A. muricata* and *K*=3 PCs for *P. damicornis*, identifying 74 and 138 outlier SNPs (q-value<0.05), respectively. The final neutral genotype matrices comprised of 5,683 SNPs for *A. muricata* and 12,815 SNPs for *P. damicornis* (**Suppl. Table S4**).

We found strong population structuring for *A. muricata*, where PCoA and sNMF models indicate the presence of three genetic clusters strongly associated with geographic distributions (**Fig. 2a**). Furthermore, we find evidence of high gene flow between reefs within regions contrasted with stronger genetic differentiation between regions. Indeed, the first two PCoA axes (capturing almost 20% of neutral genetic variation) assigned individuals into groups closely reflecting sampled regions (**Fig. 2ai**), with sNMF models confirming that individuals are genetically similar within regions with low gene flow between regions (**Fig. 2aii-iii**). These patterns are statistically supported by AMOVAs, where genomic variance is significantly higher between regions than expected by chance (*p*<0.01; **Suppl. Fig. S4a**). In contrast, variance between sites within regions and variance between subregions were not significant. Low pairwise F_ST_ values (i.e., F_ST_≤0.01) further substantiates substantial gene flow with limited genetic differentiation between reefs of the same region (Table 2). In contrast, pairwise F_ST_ values between certain regions exceeded 0.20, signalling restricted gene flow and increased genetic divergence between more distant regions (Table 2); larger F_ST_ values (i.e., F_ST_≥0.20) were between reefs at the Seychelles with Rodrigues, as well as between site S04 on the western side of Mahé (SEY) with reefs around the main islands of Mauritius. Last, high inbreeding coefficients (F_IS_>0.20) were observed between reefs of the Seychelles, Rodrigues, and the northern and southern reefs of Mauritius (**Suppl. Table S5a**), further pointing towards restricted gene flow between reefs of different regions.

**Figure 2.**
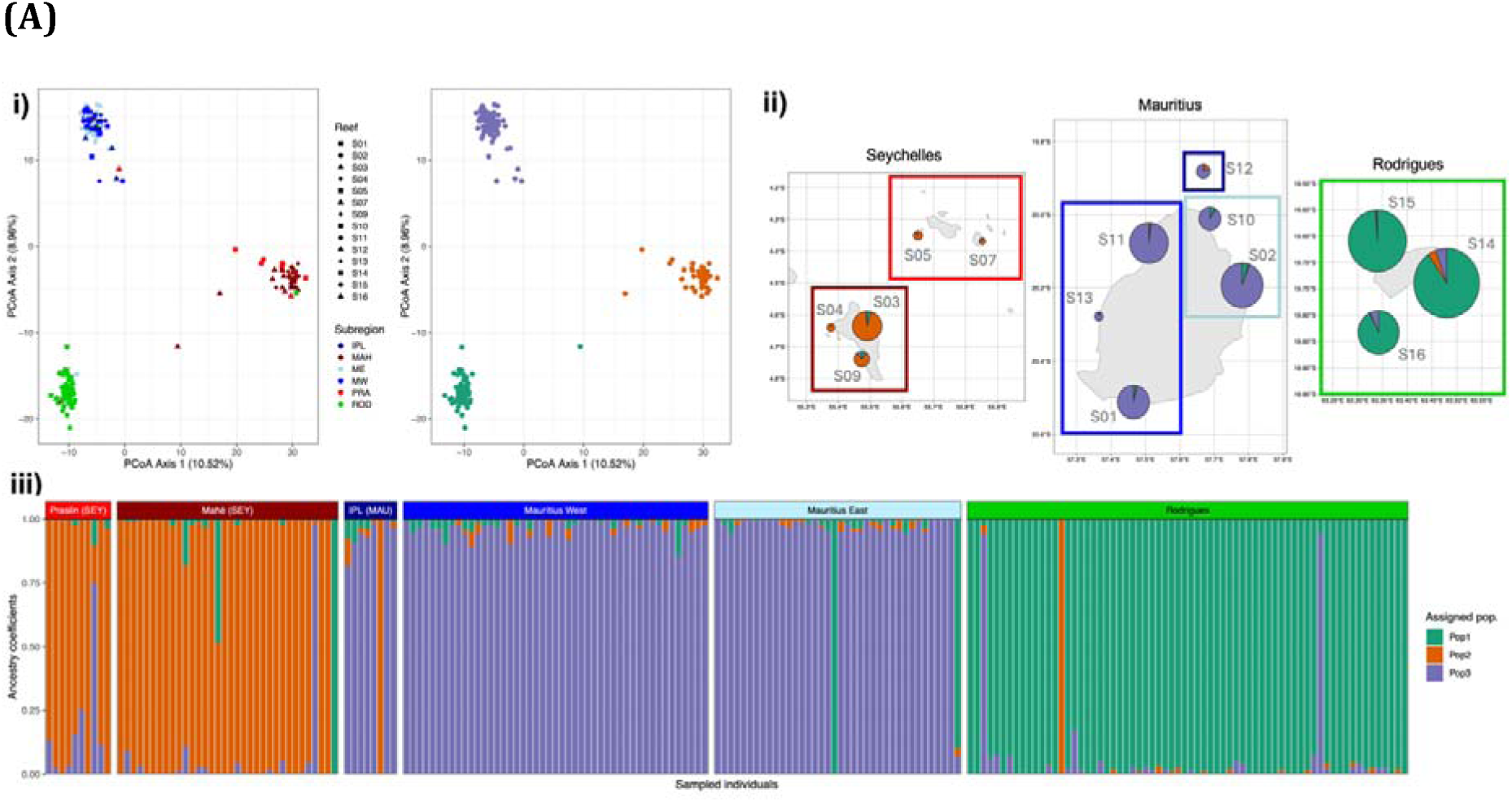
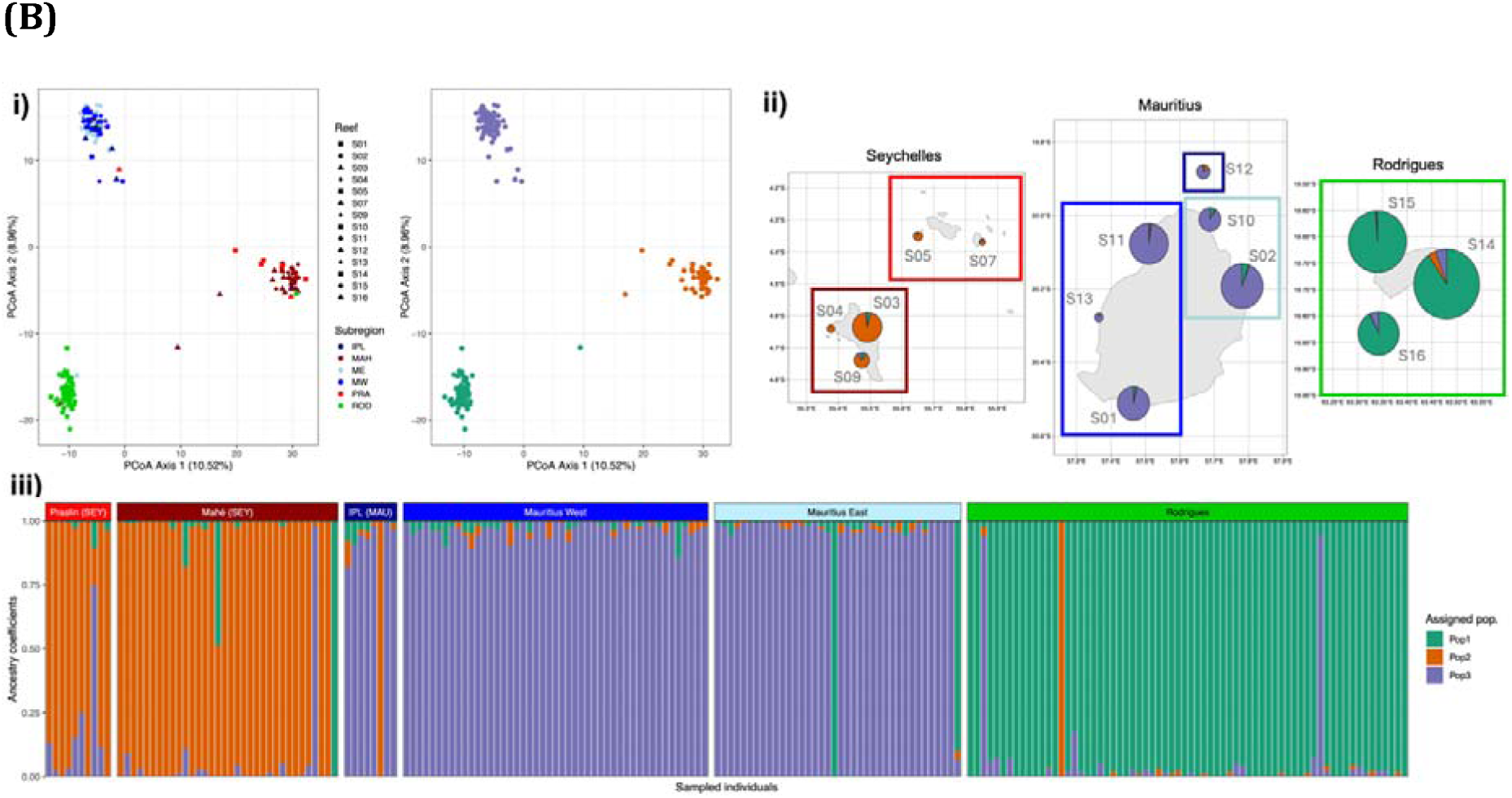
Population structure analyses of **(A)** *A. muricata* and **(B)** *P. damicornis*, assessed by a principal coordinate analysis (PCoA) and sparse nonnegative matrix factorization (sNMF), with *K*=3 and *K*=4 groups identified for each species respectively in both methods. **i)** Individual scores on PCoA axes 1 and 2 grouped by geographic location (shapes indicate reef and colours indicate subregion; left) and hierarchical cluster grouping of PCoA results (coloured by assigned grouping; right). **ii)** Maps indicating the proportion of individuals at each sampled reef allocated to *K* ancestral populations using sNMF admixture models, where pie chart size indicates the relative number of sampled individuals at each reef and coloured boxes indicate sub-regions. **iii)** Proportion of sampled individuals (columns) allocated to *K* ancestral populations using sNMF admixture models, with individuals grouped by sub-region. Sub region abbreviations: PRA = Praslin, MAH = Mahé, IPL = Ile Plate, MW = West Mauritius, ME = East Mauritius, ROD = Rodrigues.

**Table 2.** Pairwise F_ST_ between study sites for *A. muricata* (lower triangle) and *P. damicornis* (upper triangle) based on neutral SNP genotype matrices. Sites are grouped and coloured by region and subregion. F_ST_ >0.20 are highlighted in yellow.

|  |  |  | *Pocillopora damicornis* |  |  |  |  |  |  |  |  |  |  |  |  |  |  |
| --- | --- | --- | --- | --- | --- | --- | --- | --- | --- | --- | --- | --- | --- | --- | --- | --- | --- |
|  |  |  | Seychelles |  |  |  |  |  | Mauritius |  |  |  |  |  | Rodrigues |  |  |
|  |  |  | PRA |  |  | MAH |  |  | IPL | MW |  |  | ME |  | ROD |  |  |
|  |  |  | S06 | S05 | S07 | S03 | S04 | S09 | S12 | S01 | S11 | S13 | S02 | S10 | S14 | S15 | S16 |
| *Acropora muricata* | Seychelles | S06 |  | 0.02 | 0.11 | 0.03 | 0.03 | 0.04 | 0.15 | 0.14 | 0.12 | 0.17 | 0.15 | 0.16 | 0.13 | 0.15 | 0.16 |
|  |  | PRA S05 |  |  | 0.06 | 0.05 | 0.02 | 0.07 | 0.12 | 0.10 | 0.10 | 0.20 | 0.17 | 0.17 | 0.11 | 0.14 | 0.14 |
|  |  | S07 |  | 0.00 |  | 0.16 | 0.09 | 0.15 | 0.08 | 0.05 | 0.10 | 0.23 | 0.21 | 0.20 | 0.07 | 0.13 | 0.10 |
|  |  | MAH S03 |  | 0.00 | 0.01 |  | 0.05 | 0.10 | 0.20 | 0.19 | 0.17 | 0.23 | 0.20 | 0.21 | 0.18 | 0.21 | 0.22 |
|  |  | S04 |  | 0.00 | 0.01 | 0.00 |  | 0.08 | 0.14 | 0.12 | 0.12 | 0.20 | 0.17 | 0.17 | 0.11 | 0.14 | 0.14 |
|  |  | S09 |  | 0.00 | -0.02 | 0.00 | 0.00 |  | 0.15 | 0.15 | 0.13 | 0.20 | 0.17 | 0.18 | 0.14 | 0.17 | 0.19 |
|  | Mauritius | IPL S12 |  | 0.12 | 0.09 | 0.14 | 0.15 | 0.09 |  | -0.01 | 0.01 | 0.15 | 0.14 | 0.13 | 0.03 | 0.07 | 0.06 |
|  |  | S01 |  | 0.18 | 0.15 | 0.20 | 0.21 | 0.15 | 0.00 |  | 0.04 | 0.18 | 0.16 | 0.16 | 0.04 | 0.08 | 0.06 |
|  |  | MW S11 |  | 0.17 | 0.15 | 0.19 | 0.21 | 0.14 | 0.00 | 0.00 |  | 0.03 | 0.05 | 0.04 | 0.03 | 0.05 | 0.08 |
|  |  | S13 |  | 0.16 | 0.13 | 0.18 | 0.20 | 0.12 | -0.01 | 0.00 | 0.00 |  | 0.03 | 0.02 | 0.14 | 0.13 | 0.19 |
|  |  | ME S02 |  | 0.17 | 0.16 | 0.19 | 0.21 | 0.14 | 0.01 | 0.00 | 0.00 | 0.00 |  | 0.01 | 0.10 | 0.11 | 0.16 |
|  |  | S10 |  | 0.17 | 0.14 | 0.18 | 0.20 | 0.13 | 0.00 | 0.00 | 0.00 | -0.01 | 0.00 |  | 0.12 | 0.12 | 0.17 |
|  | Rodrigues | S14 |  | 0.19 | 0.17 | 0.19 | 0.22 | 0.15 | 0.10 | 0.12 | 0.12 | 0.11 | 0.11 | 0.10 |  | 0.01 | -0.01 |
|  |  | ROD S15 |  | 0.21 | 0.20 | 0.21 | 0.24 | 0.17 | 0.13 | 0.14 | 0.14 | 0.14 | 0.13 | 0.13 | 0.00 |  | 0.04 |
|  |  | S16 |  | 0.20 | 0.19 | 0.21 | 0.23 | 0.16 | 0.11 | 0.12 | 0.12 | 0.12 | 0.11 | 0.11 | 0.00 | 0.00 |  |

Sub-region abbreviations: PRA = Praslin, MAH = Mahé, IPL = Île Plate, MW = West Mauritius, ME = East Mauritius, ROD = Rodrigues

**Figure 3.**
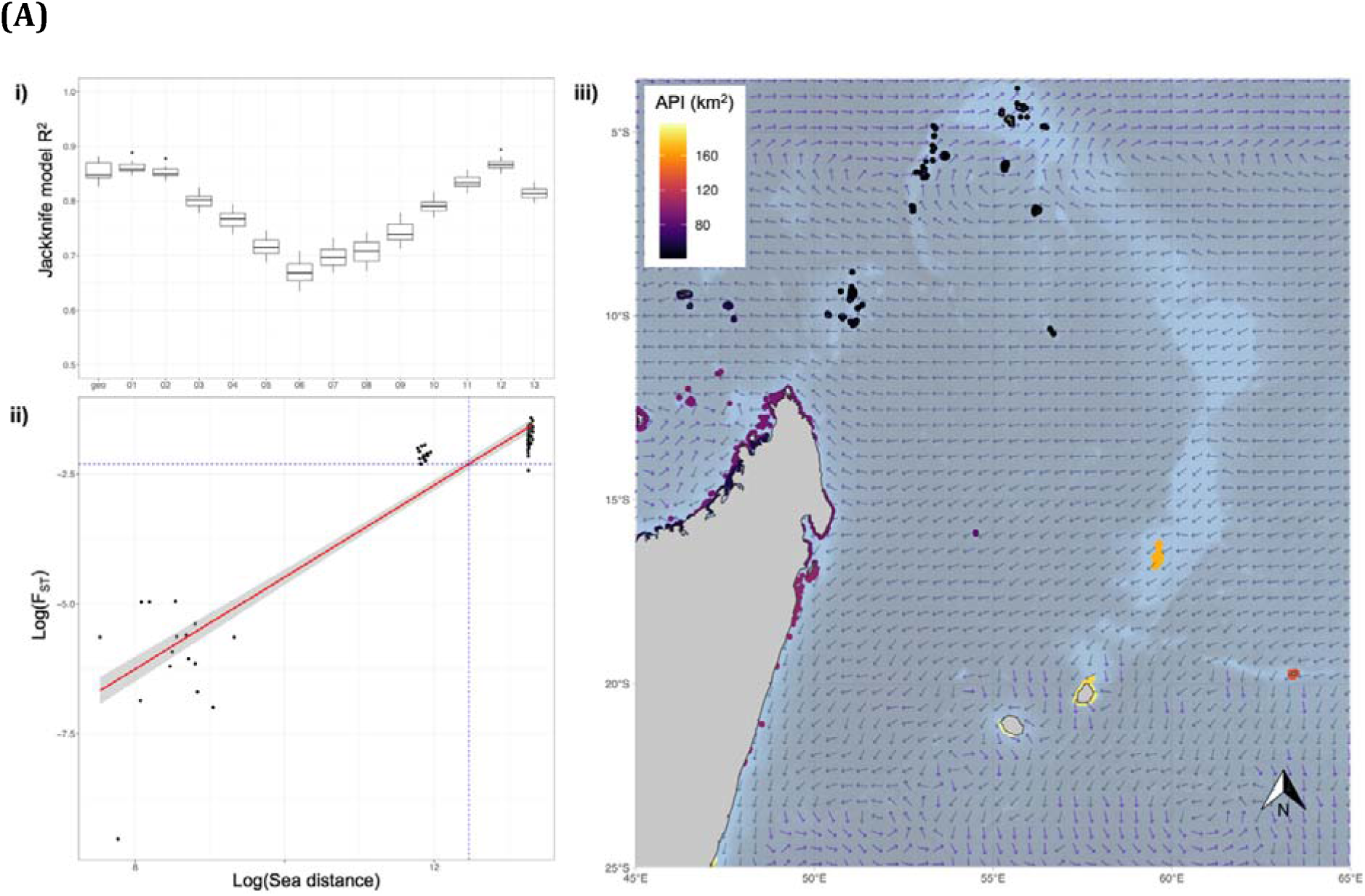
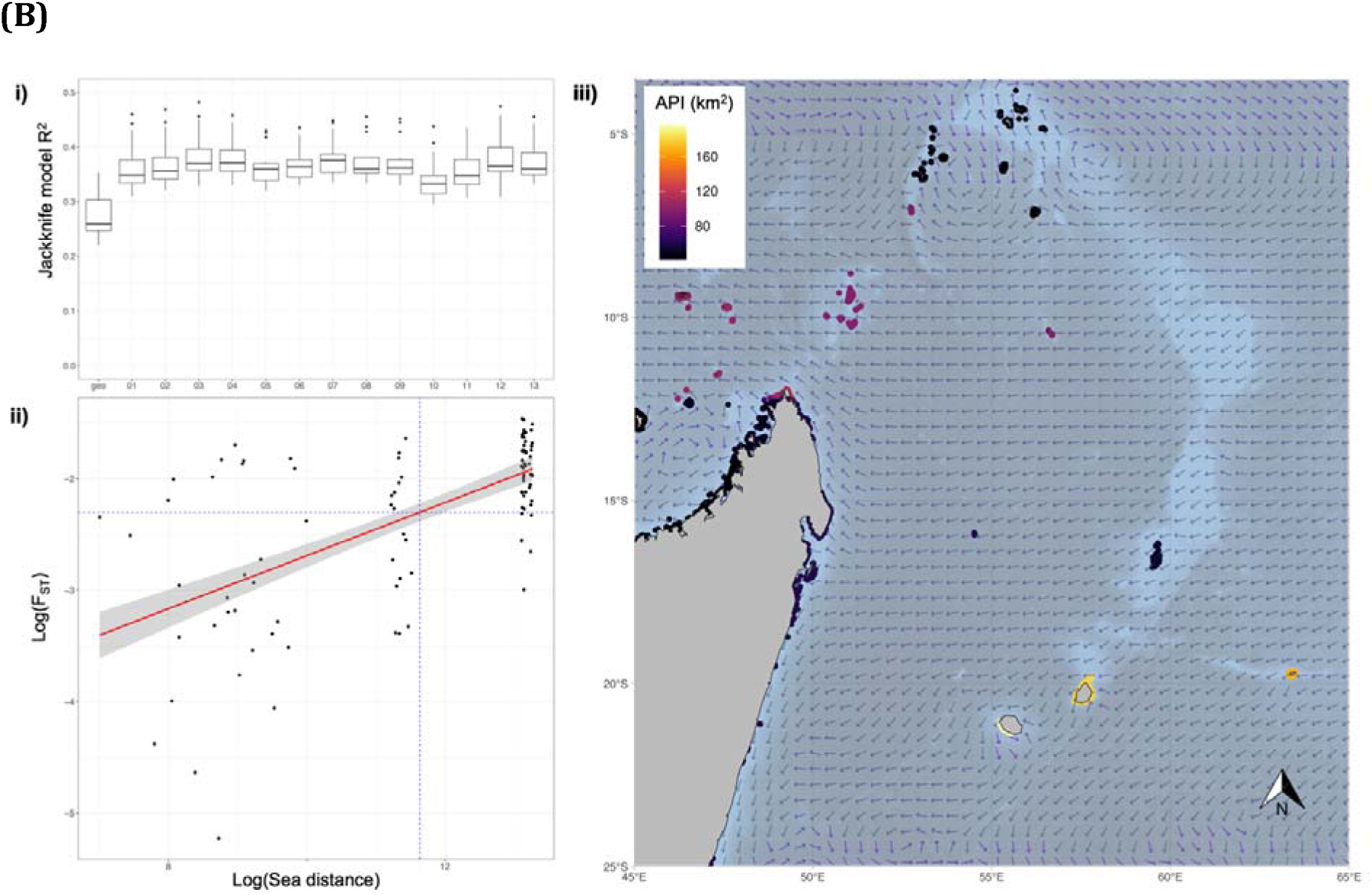
Adaptive potential indices and predictive sea currents for **(A)** *A. muricata* and **(B)** *P. damicornis*. **i)** Connectivity model performance boxplots, measured as R^2^, for models describing the relationship between log_10_ of the genetic distance (F_ST_) and log_10_ of the geographic distance, where boxplots show the spread of R^2^ values of connectivity models built with leave-one-reef-out jackknife analyses, assessing the Euclidean geographic distance (‘geo’) and sea current averaged over each month (01–12) and the year (13). **ii)** Log_10_-log_10_ relationship between the genetic distance (F_ST_) and sea distance obtained from the strongest connectivity predictor (December sea currents for *A. muricata* in **(A)**; annual sea currents for *P. damicornis* in **(B)**). The blue dotted lines indicate the sea distance that corresponds to F_ST_=0.1. **iii)** Adaptive Potential Index (API in km^2^) across the whole WIO region, calculated by combining Inbound Connectivity and Adaptive Scores. Larger API indicates more probability of hosting adaptive genotypes up-current of connected reefs. Arrows indicate sea current velocity of the strongest connectivity predictor.

**Figure 4.**
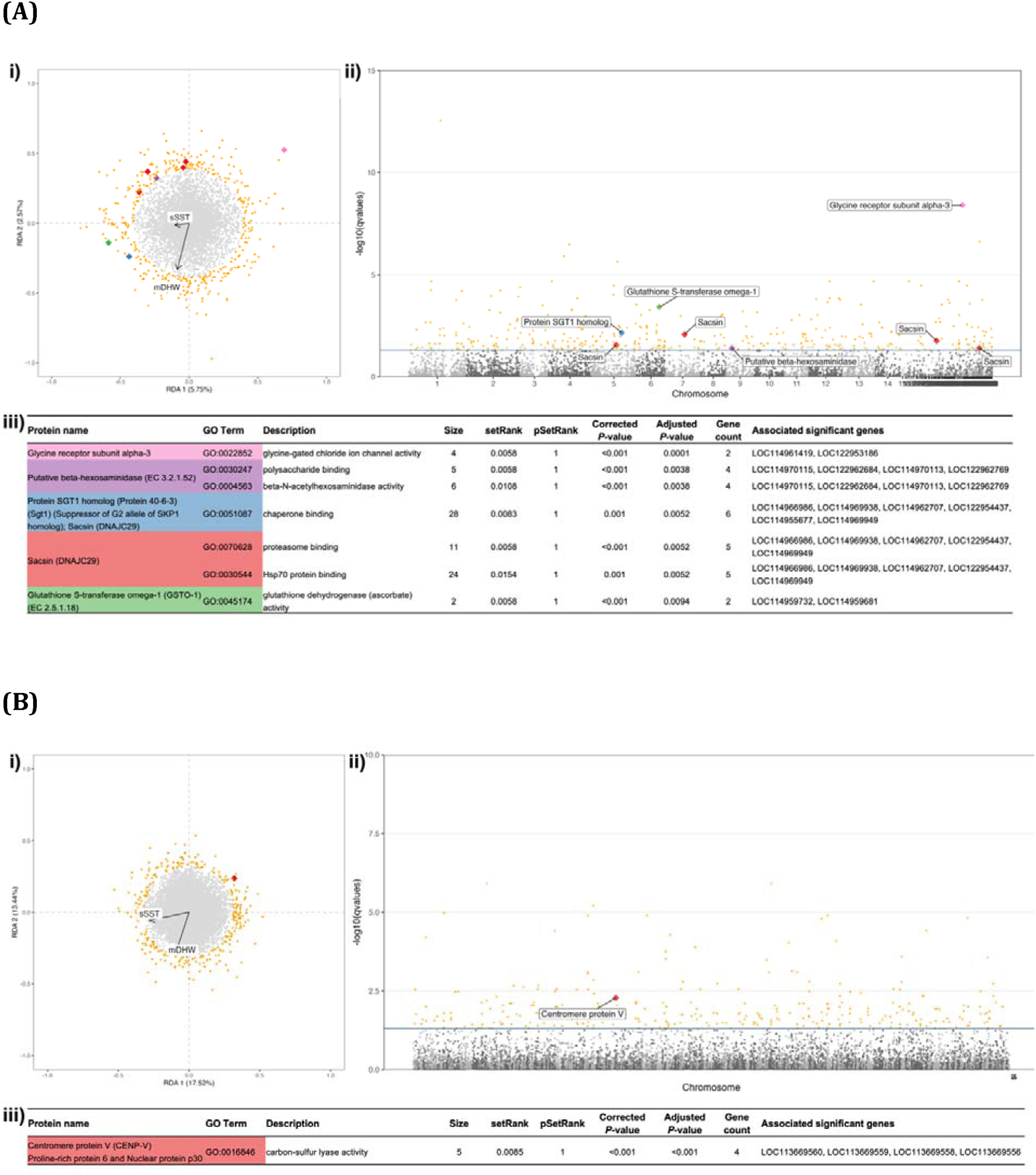
Genotype–environment association results for **(A)** *A. muricata* and **(B)** *P. damicornis*, using multivariate partial RDA (redundancy analyses) at the reef-level. **i)** RDA biplots projecting loci (dots) and environmental variables (arrows, where sSST is the standard deviation of sea surface temperature and mDHW is the mean of the degree heating weeks). Locus scores are rescaled by a factor of 12 to improve visibility. **ii)** Manhattan plots with the distribution of −log10(q-values) in the genomes. For the first two panels, loci that were identified as outliers (q- value<0.05) are coloured in orange, where outlier loci found within ±10kbp of significantly enriched genes (determined using SetRank) are labelled as diamonds and colour-coded by protein function. **iii)** Tables summarising the significantly enriched gene ontology (GO) terms and their associated protein names and descriptions, alongside SetRank results and the associated genes. Protein names and GO terms are colour coded to match panels **i)** and **ii)**.

Four genetic clusters were identified for *P. damicornis* across the three geographic regions using PCoA (first two axes capturing 23% of neutral genetic variance) and sNMF. The genetic clusters roughly matched geographic location, with evidence of some gene flow between geographic regions (**Fig. 2b**). AMOVA analyses indicate that genomic variance is significantly higher between sites (*p*<0.01) and between regions (p=0.03) than expected by chance, while variance between subregions were not significant (**Suppl. Fig. S4b**). There appears to be strong genetic similarities between individuals in the northern and southern Mauritius reefs with individuals in Rodrigues, as well as with some individuals in the northern Seychelles (Praslin) reefs (yellow clusters in **Fig. 2b**). Within regions, reefs had varying levels of genetic differentiation, with pairwise F_ST_ values between -0.01 and 0.16. Most individual genetic similarity was between the Rodrigues reefs, while the largest differences were between the Mauritius island reefs (Table 2). Between regions, the largest differences were between the eastern and western reefs of Mauritius with the Seychelles (F_ST_≥0.15), while the most similar regions are between those of the Mascarene Islands (i.e., Mauritius and Rodrigues; F_ST_<0.15). Finally, most sites had evidence of panmixia, though higher than expected homozygosity (F_IS_≥0.20) was observed at the eastern-most reef of the Seychelles (S07), as well as at the northern (S12) and southern (S01) Mauritius reefs (**Suppl. Table S5b**).

### 3.3 Connectivity Analysis

Genetic connectivity between *A. muricata* populations was well explained by sea currents and geographic distance, with a median R^2^ of 0.80 (range of median R^2^ 0.67–0.87) across all predictors following a leave-one-reef-out jackknife analysis (**Fig. 3ai**). The best predictor of F_ST_ was mean sea current in December with a median R^2^ of 0.87 (IQR 0.86–0.87; **Fig. 3ai**). Genetic connectivity between *P. damicornis* populations was also well explained by sea currents, albeit to a lesser degree than for *A. muricata*. Monthly sea currents all had similar strengths to predict patterns of genetic connectivity of *P. damicornis* across the WIO, with associations producing a median R^2^ of 0.36 (range of median R^2^ 0.26–0.38; **Fig. 3bi**). Geographic distance again explained less genetic connectivity than sea currents, with a median R^2^ of 0.26 (IQR 0.25–0.40). We therefore chose to model the genetic connectivity of *P. damicornis* using the average annual sea current, which explained 36% of the F_ST_ variation (median R^2^ of 0.36; IQR 0.35–0.39; **Fig. 3bi**).

The species-specific log_10_–log_10_ relationships between F_ST_ and optimised sea distance (based on December currents for *A. muricata* and annual currents for *P. damicornis*) indicate a stronger relationship by approximately 3.5 times for *A. muricata* than for *P. damicornis*, with respective regression coefficients of 0.885 (**Fig. 3aii**) and 0.238 (**Fig. 3bii**). These results reiterate patterns of neutral gene flow observed between regions for both species (see Section 3.2), where A*. muricata* individuals were more genetically similar between closer reefs and more genetically dissimilar with further reefs, while the weaker relationship for *P. damicornis* highlights that physical distance does not necessarily correspond with genetic similarity amongst individuals.

### 3.4 Genotype–Environment Associations to identify loci under selection

The two thermal stress variables, sSST and mDHW, best explained reef-level neutral genetic variation for both *A. muricata* and *P. damicornis*, as determined using an automated forward and backward variable selection method. As a preliminary partial RDA revealed confounding effects of neutral genetic structure and thermal stress, we performed GEA analyses while accounting for neutral structure using 3 PCs for *A. muricata* (explaining 68.4% variance) and 4 PCs for *P. damicornis* (explaining 64.2% variance) to reduce false positive outlier loci.

For *A. muricata*, the first and second pRDA axes respectively explained 21.0% and 5.8% of genetic variation (**Fig. 4ai**). Both axes were negatively associated with sSST and mDHW, with the second axis showing a stronger association with mDHW. A total of 380 outlier loci were identified (*q*-value<0.05) throughout the genome (**Fig. 3aii**). SetRank analyses revealed that significant genes (i.e., those within ±10 kbp of significant SNPs) were associated with seven enriched GO terms (molecular functions) and encode five proteins: glycine receptor subunit alpha-3 (GO:0022852), putative beta-hexosaminidase (GO:0030247, GO:0004563), protein SGT1 homolog (GO:0051087), Sacsin (GO:0070628, GO:0030544) and Glutathione S- transferase omega-1 (GO: 0045174) (**Fig. 4aiii**). Interestingly, six genes significantly associated with temperature were annotated as Sacsin (DNAJC29) and SGT1 homolog proteins, which have the molecular function “chaperone binding activity” (of these, five genes were also associated with “Hsp70 protein binding activity”). The other significant genes included those associated with glycine-gated chloride ion channel activity (two genes), polysaccharide binding activity (specifically with beta-N-acetylhexosaminidase activity; four genes), and glutathione dehydrogenase activity (two genes).

In *P. damicornis*, the first pRDA axis explained 17.8% of genetic variation and was negatively associated with sSST, while the second partial RDA axis explained 16.9% of genetic variation and was negatively associated with mDHW (**Fig. 4bi**). A total of 292 outlier loci were identified (**Fig. 3bii**). SetRank enrichment analyses identified only carbon-sulfur lyase activity (GO:0016846) as an enriched molecular function by four significant genes that are associated with centromere protein V (**Fig. 4biii**).

### 3.5 Adaptive Potential Index (API) across the West Indian Ocean

We combined network-level connectivity models with spatial distributions of putatively heat- adapted genotypes to produce an Adaptive Potential Index (API) across the WIO for identifying donor reefs with the potential to disperse thermal-adapted genotypes to neighbouring reefs (Table 3). For a given pixel, the API represents the surface area (in km²) of neighbouring reefs that (1) have a thermal history linked to the presence of candidate adaptive loci (Adaptive Score ≥0.8) and (2) are within a sea distance indicative of inbound connectivity (i.e., a distance that corresponds to an F_ST_ ≤0.1 upstream of the target reef). For both species, Adaptive Scores (AS) and API were region-specific, with higher values in more southern reefs at Mauritius and Rodrigues and lower values at reefs near the equator at the Seychelles (**Fig. 3iii**; Table 3; **Supp Fig. S5**).

**Table 3.**
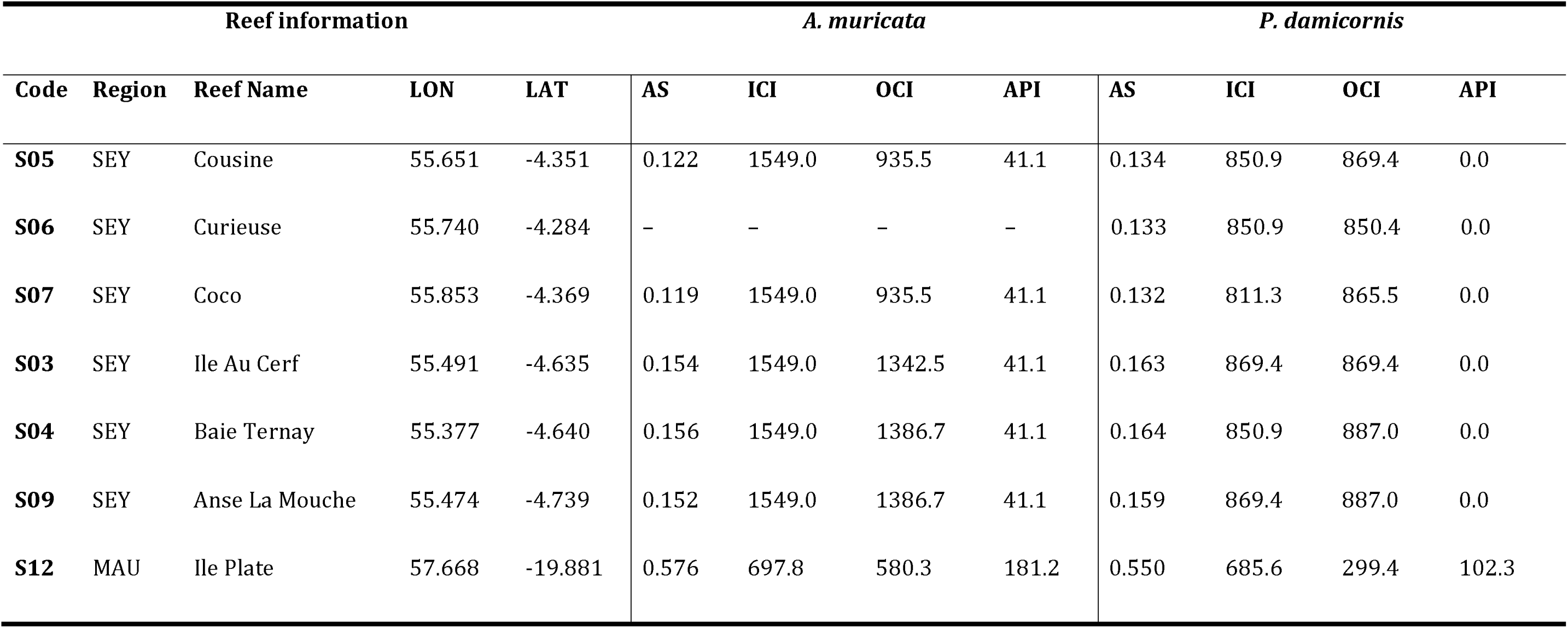
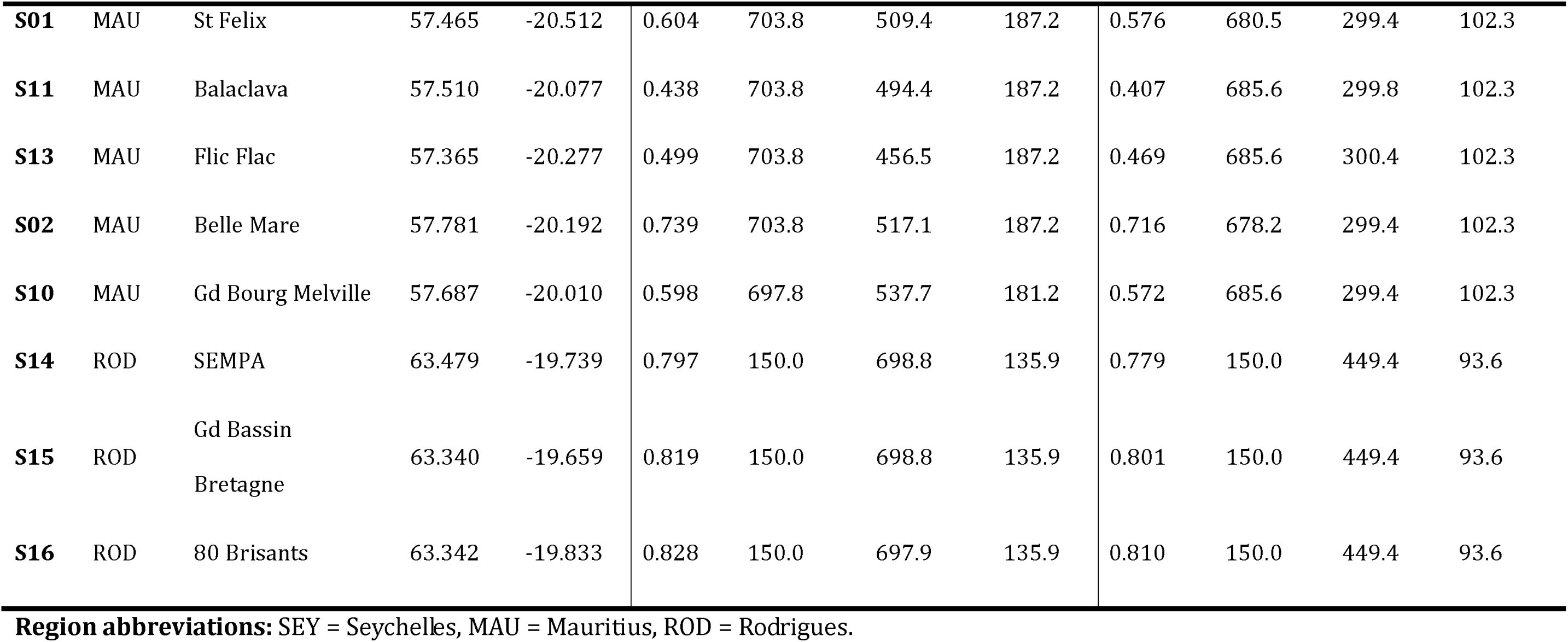
Adaptive potentials and connectivity metrics for *A. muricata* and *P. damicornis* at 15 coral reefs across the West Indian Ocean. The Adaptive Score (AS) indicates the relative level of adaptation to thermal stressors (sSST and mDHW) from 0 to 1 (low to high adaptive values), as determined by a GEA analysis. Inbound and Outbound Connectivity indices (ICI and OCI, respectively) indicate the respective area (in km^2^) of reef up-current and down-current that fall within a sea distance corresponding to genetic connectivity of F_ST_<0.1. The Adaptive Potential Index (API) indicates the area (in km^2^) of reefs upstream (ICI) that contain adapted corals (AS>0.8). See **Supp Fig. S5** for spatial representations.

For *A. muricata* (**Fig. 3aiii**; Table 3), the reefs around Mauritius hosted moderately heat adapted corals (AS between 0.43–0.74), further receiving adapted propagules from an estimated 181–187 km^2^ of other reefs. While *A. muricata* at the Rodrigues reefs had slightly reduced API values (136 km^2^) due to fewer heat adapted propagules up-current, these corals had the highest Adaptive Scores (AS ≥0.80) of the studied reefs, supplying propagules to almost 700 km^2^ of reefs downstream (OCI). In the northern WIO, *A. muricata* corals of the Seychelles had both lower thermal Adaptive Scores (AS <0.16) and the lowest API with heat adapted propagules coming from an estimated 41 km^2^ of reefs (API), despite these reefs being connected to approximately 1550 km^2^ of reefs upstream (ICI) and >935 km^2^ downstream (OCI).

Similar AS estimates of coral adaptation to thermal stress and patterns of API were observed for *P. damicornis*, where connectivity measurements based on weaker F_ST_ relationships with sea currents resulted in lower values for IBI, OCI and API (**Fig. 3biii**; Table 3). The reefs at Mauritius and Rodrigues appear to host the most heat adapted corals (AS between 0.41–0.81), respectively receiving heat adapted propagules from an estimated 102 km^2^ and 94 km^2^ of reef (API) and providing adapted propagules to approximately 300 km^2^ and 450 km^2^ of reef (OCI). In the Seychelles, *P. damicornis* corals had very low adaptive scores (AS <0.17) and were expected to receive no adapted propagules (API of 0 km^2^) from the 811–870 km^2^ of inbound connected reefs.

## 4. Discussion

### 4.1 Life-history traits and asymmetric ocean currents drive population structure across the WIO

Coupling genomic information with environmental data revealed the importance of coral life-history strategies and asymmetrical ocean currents in driving spatial patterns of population structure across the WIO. The population structure uncovered in this study corroborates connectivity simulations and mapped genetic structuring of various other coral species across the WIO (e.g., Gélin et al., 2018; Kusumoto et al., 2020; Oury et al., 2024; Vogt-Vincent et al., 2024).

*A. muricata* showed strong patterns of local genetic mixing with minimal between-region gene flow across the WIO, mirroring expectations for a synchronous broadcast spawning coral species (Miller and Ayre, 2008). Indeed, high gene flow between reefs of a region within a relatively short distance has been found for numerous Acroporidae around the world (e.g., Adam et al., 2022; Fuller et al., 2020; Lukoschek et al., 2016; Nakajima et al., 2010; Selmoni et al., 2021). Genetic mixing is promoted by synchronous release of gametes (Wijayanti et al., 2019), while a free-swimming larval stage allows propagule dispersal to neighbouring reefs (Tabitha et al., 2021). Some gene flow occurs between southern reefs of the Mascarene Islands (Mauritius and Rodrigues) in a predominately westward direction along ocean currents (Vogt-Vincent et al., 2024). However, corals in the Seychelles appear to be genetically isolated, where gene flow in the Austral summer during *A. muricata* spawning is blocked by the strong oceanographic barrier of the South Equatorial Countercurrent (at approximate latitude of 10° S; Phillips et al., 2021).

Similar, albeit more complex, population structuring was found for *P. damicornis* with three region-specific genetic clusters and a fourth dispersed genetic cluster (yellow group in **Fig. 2b**). The general pattern of gene flow between the Mascarene Islands reflects that observed for the *A. muricata* populations and that of previous investigations into *P. damicornis* Type β dispersal across the southern WIO (Gélin et al., 2018). The genetic dissimilarity observed for our *P. damicornis* between the northern and southern regions can again be explained by strong ocean currents (Phillips et al., 2021). The fourth genetic cluster poses an interesting complexity, potentially attributed to contemporary gene flow, remnants of a common ancestral population, and/or the inclusion of cryptic species or ecotypes in our analyses. Contemporary region-wide gene flow might be facilitated by a well provisioned larval stage with up to 14 weeks viability (Gélin et al., 2018, 2017; Richmond, 1987), where propagules can survive long distances, travelling passively by ocean currents (Phillips et al., 2021), rafting (Nikula et al., 2013), or in boat ballast water or hull fouling (Gollasch, 2008). Alternatively, this anomalous genetic cluster could represent genetic remnants of a common ancestral population (Oury et al., 2024) that expanded across the WIO following the penultimate glacial period (ca. 100,000 years ago) under altered sea current conditions (Colleoni et al., 2016; Herbert et al., 2010). Evidence of ancestral genetic make-up could be preserved by high rates of selfing and asexual reproduction of *P. damicornis* (indeed, 65% of our samples were clones). Finally, the genetic cluster could simply represent the inclusion of hybridised individuals, either between closely related cryptic species occurring in sympatry (Carr et al., 2025; Combosch and Vollmer, 2015; Gélin et al., 2018; Oury et al., 2024) or between colonies of genetically divergent and locally adapted ecotypes (e.g., shallow lagoon ecotypes mating with outer reef slope ecotypes; Oury et al., 2024).

Regardless of the mechanism, population structure is rarely a simple function of reproductive mode or larval type (Miller and Ayre, 2008; Prata et al., 2024; Severance and Karl, 2006). While genetic exchange seems to occur across the southern WIO from Rodrigues towards the other Mascarene Island reefs, and potentially beyond to Madagascar and the African east coast, the South Equatorial Countercurrent barrier is undoubtedly inhibiting gene flow between southern to northern WIO reefs near the equator (e.g., Seychelles). This can have important consequences for adapted genotype exchange across the WIO, with impacts on management strategies to be implemented in different regions (Phillips et al., 2021; Vogt-Vincent et al., 2024). More wide-spread genetic sampling of corals is required to draw precise conclusions of population structure across the WIO region.

### 4.2 Sea currents as a proxy for genetic connectivity and dispersal

Using sea currents to determine asymmetric distance between reefs was a better predictor of population connectivity than Euclidean geographic distance for both coral species, supporting the growing body of literature calculating gene flow for marine organisms (e.g, Benestan et al., 2016; Cooke et al., 2016; Hernawan et al., 2023; Krueck et al., 2020; Prata et al., 2024; Snead et al., 2023; Villastrigo et al., 2023; White et al., 2010). For synchronous broadcast spawning corals, we might expect that genetic distance would be best predicted using sea currents from a particular time of the year. Indeed, connectivity of *A. muricata* was best predicted by sea currents in December, reflecting timing of known mass spawning for this species (Wijayanti et al., 2019). When no month’s sea currents clearly explain genetic distance, such as for *P. damicornis*, it may be that the species has asynchronous spawning and potentially complex reproductive modes.

The correlation (r^2^) and regression coefficients (m) of the connectivity models can also provide information about the pattern and strength of sea currents used to disperse the propagules across a reef network. A strong correlation coefficient seems to indicate more local recruitment of propagules, such as for *A. muricata* (r^2^ =0.87), while weaker correlation coefficients indicate decoupling of gene flow from physical geographic distance, such as for *P. damicornis* (r^2^ =0.37). The regression coefficient provides a rough estimate of length of larval stage, where the 3.5 times steeper regression coefficient of *A. muricata* vs *P. damicornis* (m =0.88 vs. m =0.24) mirrors the much shorter larval stage (3 weeks vs 14 weeks; Richmond, 1987; Tabitha et al., 2021). By using metrics derived from the connectivity model, hypotheses of life history strategies can be obtained for non-model coral species to facilitate the development of appropriate management strategies.

### 4.3 Thermal stress drives signals of local adaptation

Historical thermal variation was found to correlate with non-neutral genetic diversity of two coral species across the WIO. Multivariate genotype–environment associations (GEAs) suggest local adaptation to thermal stress, with 380 and 292 outlier SNPs for *A. muricata* and *P. damicornis*, respectively, significantly associated with standard deviations in sea surface temperatures (sSST ; Skirving et al., 2020) and mean degree heating weeks (mDHW; Skirving et al., 2020). In both species, corals from Rodrigues exhibited the strongest signals of thermal adaptation, likely due to the higher and more fluctuating temperatures of its lagoon and reef flat environments. In contrast, corals from the Seychelles showed the weakest adaptive signals, as their backreef and reef slope habitats are influenced by cooler upwelling waters. These findings reflect recent evidence of mean SST as a strong predictor of global genetic diversity in *P. damicornis*, with highest nucleotide diversity at reefs experiencing higher temperatures (Carr et al., 2025).

Extreme and variable marine thermal conditions can trigger coral bleaching when SST fluctuations exceed seasonal averages (Hughes et al., 2017b; Liu et al., 2003), a pattern observed across the Indian Ocean (Shlesinger and van Woesik, 2023). Recurrent mass-bleaching events over decades exert strong selective pressures (Hughes et al., 2017b; Skirving et al., 2020), likely driving molecular signatures of thermal adaptation and tolerance in affected populations (Lachs et al., 2023). SST variation and mean DHW capture both the duration and intensity of thermal anomalies (Humanes et al., 2022; Liu et al., 2003; Skirving et al., 2020; van Hooidonk et al., 2016), which may explain the high number of outlier loci associated with heat stress detected here. While environmental data were extracted at 5 km resolutions, we acknowledge that reef systems can exhibit substantial microhabitat variation at much finer spatial scales (e.g., <100m), potentially influencing highly localised patterns of coral adaptation (e.g., Bay and Palumbi, 2014; Bozec et al., 2025). To mitigate this effect, we standardised sampling depth between sites and used variables from the Allen Coral Atlas (at 10m resolution) to refine the ecological characterisation of each sample site. Nonetheless, the potential for undetected fine-scale environmental heterogeneity remains a limitation here.

### 4.4 Candidate molecular targets of heat stress

Genes near SNPs significantly associated with historical heat stress exposure (sSST and mDHW) had molecular functions that were previously observed in coral thermal responses. Interestingly, we found significant enrichment for heat shock proteins and molecular chaperones in *A. muricata*, driven by the presence of five significant genes encoding the Sacsin protein. Sacsin is a co-chaperone for the heat-shock protein Hsp70 (Ménade et al., 2018; Parfitt et al., 2009) and is significantly up-regulated when *Acropora* and *Pocillopora* species are exposed to high temperatures in lab conditions (Cunning et al., 2018; Cunning and Baker, 2014; Hemond et al., 2014; Mayfield et al., 2018). Sacsin has previously been identified as a key candidate protein involved in thermal responses of *A. millepora* in the Great Barrier Reef (Fuller et al., 2020), while another seascape genomic study on *A. millepora* in New Caledonia found significant enrichment of chaperone binding proteins associated with bleaching alert frequency (Selmoni et al., 2021).

Three other molecular functions were significantly enriched for *A. muricata* in association with thermal stress. The glutathione S-transferase omega-1 (EC 2.5.1.18) enzyme was detected with two significant genes associated with sSST and mDHW. This enzyme is upregulated in some coral species following long-term (60 day) heat stress exposure (Dias et al., 2019). This enzyme plays a key role in coral cellular antioxidant and immune defence responses (Morrow et al., 2012), likely responding to reactive oxygen species induced by enhanced metabolic rates (Burdon et al., 1990; Richier et al., 2006). We also found enrichment for the lysosomal enzyme beta-hexosaminidase (EC 3.2.1.52) from four genes. This is an important factor involved in a range of functions, including glycosylation of proteins, cellular signalling, protein folding, and mycobacterium defence (Koo et al., 2008; Lundgren et al., 2013; Sproles et al., 2019). As stressed corals have increased susceptibility to disease (Littman et al., 2010; Mydlarz et al., 2010; Thurber et al., 2009), an adaptive increase in immune responses due to increased temperatures might be expected (Lundgren et al., 2013). Last, the glycine receptor (subunit alpha-3) protein associated with glycine-gated chloride ion channel activity was detected by two genes, which is involved in synaptic signalling of the central nervous system of vertebrates (Ceder et al., 2024) that might alter mobility and movement for feeding in heat-adapted corals.

Among heat-associated SNPs in *P. damicornis*, only one molecular function was significantly enriched: carbon-sulphur lyase activity. To our knowledge, this function has not been previously associated with heat stress and may represent a false signal. Given that previous studies on *Pocillopora* spp. identified candidate genes involved in immunity, cellular homeostasis, metabolism, and signalling responses to environmental stress (Lundgren et al., 2013; Mayfield et al., 2018; Poquita-Du et al., 2024; Selmoni et al., 2021), we expected a greater number of significant functional associations. The limited detection of enriched functions may be attributed to the strong population structure observed, which we attempted to control for by including neutral genetic structure in our RDA analysis. In this way, we targeted shared genomic pathways involved in thermal stress responses amongst diverged individuals, where we cannot rule out the retention cryptic or hybrid individuals in our dataset that inhabit different niches and express different responses to stressors (Johnston et al., 2017). Alternatively, a lack of signal may be attributed to the high proportion of unique, predominantly unannotated genes in *P. damicornis*, which comprise up to a quarter of its genome (Cunning et al., 2018).

We stress caution in over-interpreting results of GEA and GO term analyses. These models generate hypotheses to facilitate understandings of environmental influence on genotypes (Lasky et al., 2023; Lotterhos, 2023; Selmoni et al., 2024). Candidate genes should therefore be treated as preliminary results requiring experimental or in-field validations (reviewed in Selmoni et al., 2024). Furthermore, numerous analytical factors can affect GEA results. Here, we traded GEA power with reduced false positive rates when we corrected for population structure (Forester et al., 2018; Lotterhos, 2023). GEAs also have reduced power to detect quantitative trait nucleotides (Lotterhos, 2023); because polygenic variation is expected to drive coral thermal adaptation (Rose et al., 2018), important genomic regions under selection are likely missed. Additional methods, such as genome-wide association studies (GWAS) or quantitative trait locus (QTL) mapping, could help strengthen conclusions drawn from seascape genomic studies (Lasky et al., 2023; Lotterhos, 2023), as could the inclusion of other genomic variants that are expected to play a major role in facilitating adaptation (e.g., structural variants; Mérot et al., 2020; Pokrovac and Pezer, 2022; Wellenreuther et al., 2019). Nevertheless, our results provide valuable hypotheses of genomic regions conserved across populations in response to thermal pressures across a discontinuous reef system to guide future coral research and assist regional coral reef management.

### 4.5 Local-level reef management is recommended across the West Indian Ocean

Coral reef ecosystems across the WIO are vulnerable to collapse under projected climatic conditions (Obura et al., 2022). As local scale stressors can affect regional level community structure (Green et al., 1987), we echo recommendations for coordinated local level management across the WIO to maintain overall coral reef health (Obura et al., 2022; Stefanoudis et al., 2023). We found that the large distances between reefs reduced the importance of regional gene flow for facilitating thermal adaptation. Indeed, the reef regions investigated represent two independent ecoregions (Obura, 2012; Spalding et al., 2007).

Furthermore, coral propagule dispersal simulations across the WIO estimates that there are over 100 generational steps of separation (about 500 years) between the Mascarene Islands and the Seychelles (Vogt-Vincent et al., 2024). As congruent spatial pattens of the adaptive potential index (API) were observed for both species despite differences in life-history traits, demography and adaptation, we recommend similar management strategies for both species.

Understandings of source locations of thermally adapted corals and connectivity with other reefs in the region can help guide conservation planning (Bozec et al., 2025; Selmoni et al., 2024, 2020a). For instance, reefs hosting corals with higher adaptive potentials could be placed in a marine protected area (Abelson et al., 2016; De Clippele et al., 2023), while reefs with low adaptive potentials might require assisted gene flow from adapted reefs (Baums, 2008; Hagedorn et al., 2021; Van Oppen et al., 2015). The Mascarene Islands are important sources of heat adapted corals due to exposure to higher temperatures with large daily fluctuations across reef lagoons and flats. As the Mascarene Islands are located up-current of Madagascar and the east African coast along the South Equatorial Current (Phillips et al., 2021; Vogt-Vincent et al., 2024), these reefs may serve as important sources of thermal resilience for west WIO populations (Matz et al., 2020; Oury et al., 2024). In contrast, the Seychelles Archipelago appears to host fewer heat adapted corals, likely due to proximity with deep oceans and upwelling that bring cooler and more consistent water temperatures (Shlesinger and van Woesik, 2023). With their relative isolation, coral reefs of the Seychelles may require assisted gene flow, importing propagules from heat adapted reefs such as east African reefs that have already experienced recurrent mass bleaching (Elma et al., 2023).

Going forward, insights obtained from these seascape genomic analyses can be integrated into management frameworks that must be co-developed with local stakeholders (Stefanoudis et al., 2023). This is important to facilitate collaboration between local managers across the regional network to standardise research and management methods, share resources and undertake research that is directly relevant to making decisions for the overall health of the coral reef ecosystem in the WIO. Extrapolations of genetic-based adaptive indices to unsampled regions can be biased, particularly if predictor variables covary differently in training vs extrapolated reefs (Yates et al., 2018), or if genetic variation is not well distributed across the environmental landscape (Capblancq and Forester, 2021). Future work will need to include seascape genomic analyses across the larger WIO network prior to making more definitive conclusions and preparing management strategies.

### 4.6 Conclusions

Local thermal conditions seem to drive thermal tolerance across reefs of the West Indian Ocean. We found strong molecular signals of heat stress adaptation in corals from thermally variable environments, such as at the Rodrigues and Mauritius Islands. The genetic variants potentially implicated in thermal adaptation were associated with genes coding for proteins implemented in heat stress response, notably the heat-shock protein co-chaperone Sacsin. However, large sea distances and strong oceanographic barriers inhibit the genetic exchange of adapted genotypes across the WIO, calling for local-level management of reefs within a framework of a larger regional management plan. Further research into phenotypic expressions of hypothesised target genes will need to be carried out to validate adaptive potentials of corals during stressed conditions. Insights from this research contribute to a growing understanding of coral adaptation to thermal stress that can inform conservation strategies aimed at preserving thermally tolerant genotypes across the WIO in the face of ongoing climate change.

## Supporting information

Suppl. Methods: Selection of sample sites

## Acknowledgements

This study was conducted with funding provided under the Coral Restoration Project (PIMS 5736) by the Adaptation Fund and implemented by the United Nations Development Programme Mauritius and Seychelles. We gratefully acknowledge the contributions of the following institutions to field sampling efforts: Albion Fisheries Research Center, Eco-Sud, Reef Conservation, Marine Conservation Society Seychelles, Mauritius Oceanographic Institute, Nature Seychelles, Seychelles Parks and Gardens Authority, and Shoals Rodrigues. We also thank the United Nations Development Programme Mauritius and Seychelles for their support in coordinating activities throughout the project.

## Data availability statement

Data for this study will soon be made available on Dryad.

