## Supplementary material for "Coral genetic structure in the Western Indian Ocean mirrors ocean circulation and thermal stress": Suppl.

**Suppl. Methods.** Selection of sample sites. *(separate document)*

**Suppl. Fig. S1**. Workflow

**Suppl. Fig. S2**. Environmental variable correlations

**Suppl. Fig. S3.** LD pruning

**Suppl. Fig. S4.** AMOVA histograms

**Suppl. Fig. S5.** Adaptive and connectivity indices

**Suppl. Table S1.** Sample site location and sample size

**Suppl. Table S2.** Environmental variable descriptions

**Suppl. Table S3.** Species genetic ID

**Suppl. Table S4.** Genomic filtering

**Suppl. Table S5.** Population statistics

**Suppl. Fig. S1.** **Workflow.** Schema of the methods, highlighting the input data and analyses conducted, where we note where in the methods sections more details can be found.

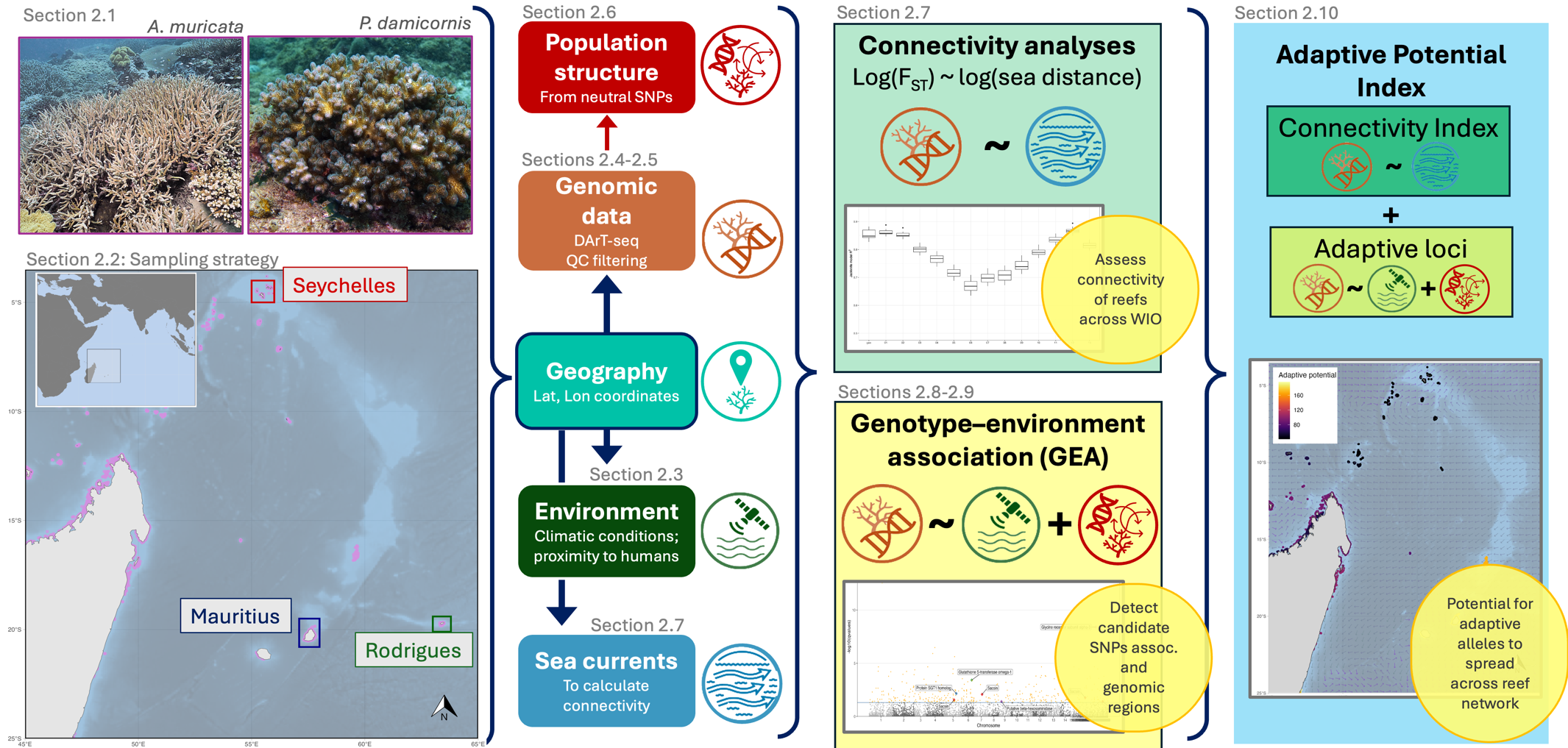

**Coral image credits:** (left) *A. muricata* taken in the Seychelles Islands by Charlie Veron; (right) *P. damicornis* taken in Brunei by Emre Turak and Lyndon DeVantier. Obtained from Corals of the World website

**Suppl. Fig. S2.** **Environmental variable correlations.** Correlation plots using Spearman non-parametric tests for (**a**) all 17 environmental variables tested and (**b**) the final 10 uncorrelated variables. See **Suppl. Table S1** for details on environmental variables and corresponding acronyms.

(**a**)

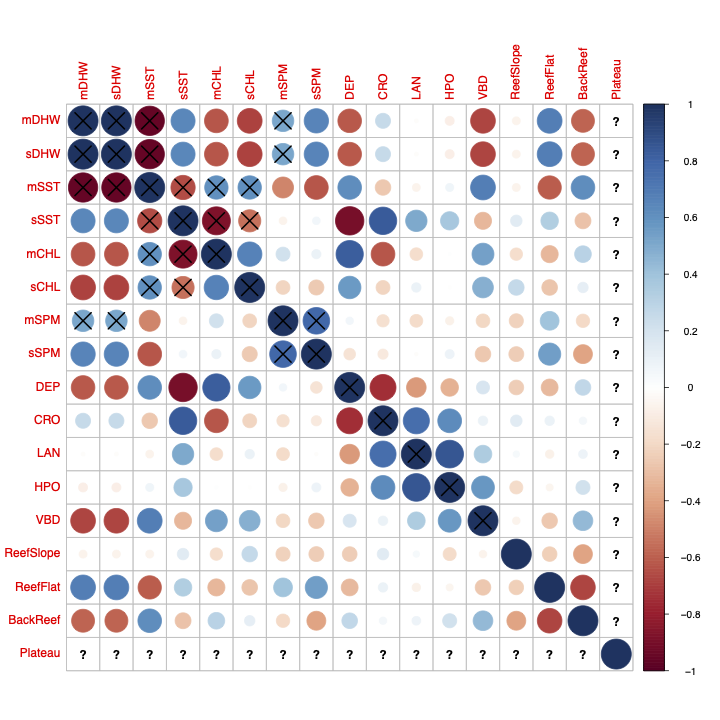

(**b**)

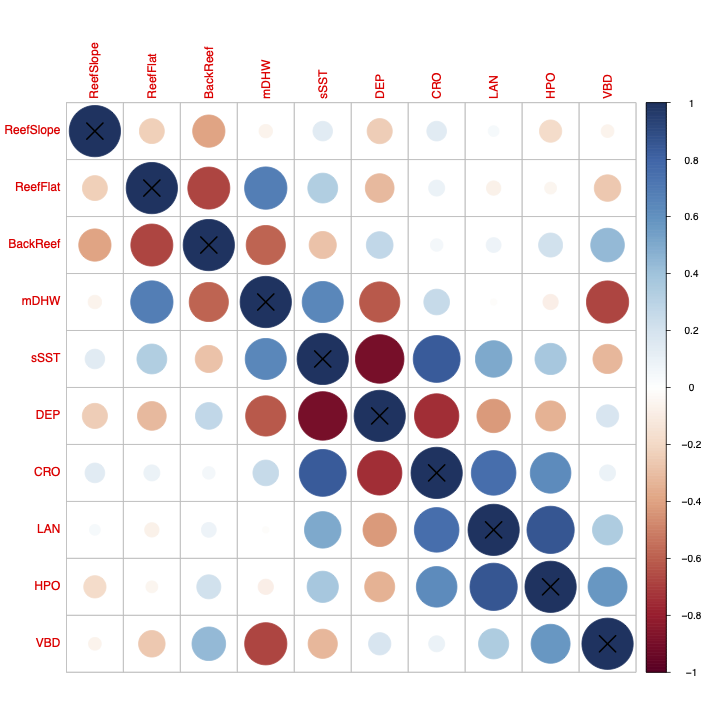

**Suppl. Fig. S3. LD pruning.** Analysis of F_ST_ and principal coordinates (PCoA) for (**a**) *Acropora* *muricata* and (**b**) *Pocillopora damicornis* pre-filtering (left column) and post-filtering (right column), including the removal of individuals identified as cryptic species. **(i)** Pre-filtering and **(ii)** post-filtering pairwise F_ST_ values, represented as heatmaps for each PCoA group with low F_ST_ in dark green and high F_ST_ in dark red. **(iii)** Pre-filtering and **(iv)** post-filtering PCoA biplots for axes 1–4, indicating the distribution of individuals in relation to each other, coded by study sites (top row, coloured by study region and shaped by site) and by PCoA-based clustering (bottom row, colours reflect different groups).

**(a) *Acropora muricata***

| **Pre-filtering** | **Post-filtering** | |
| --- | --- | --- |
| **i)**  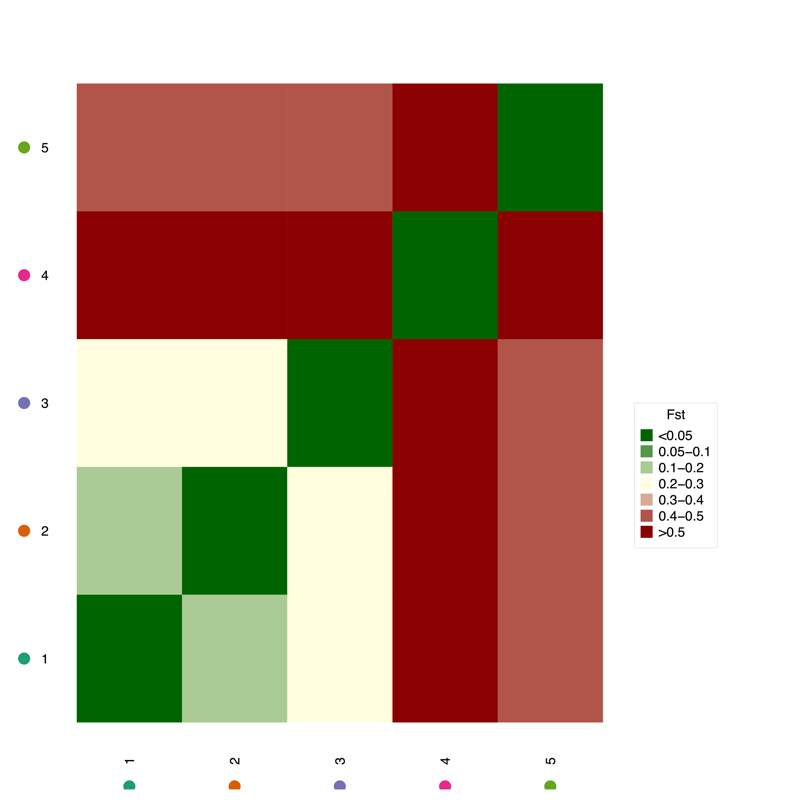 | | **ii)**  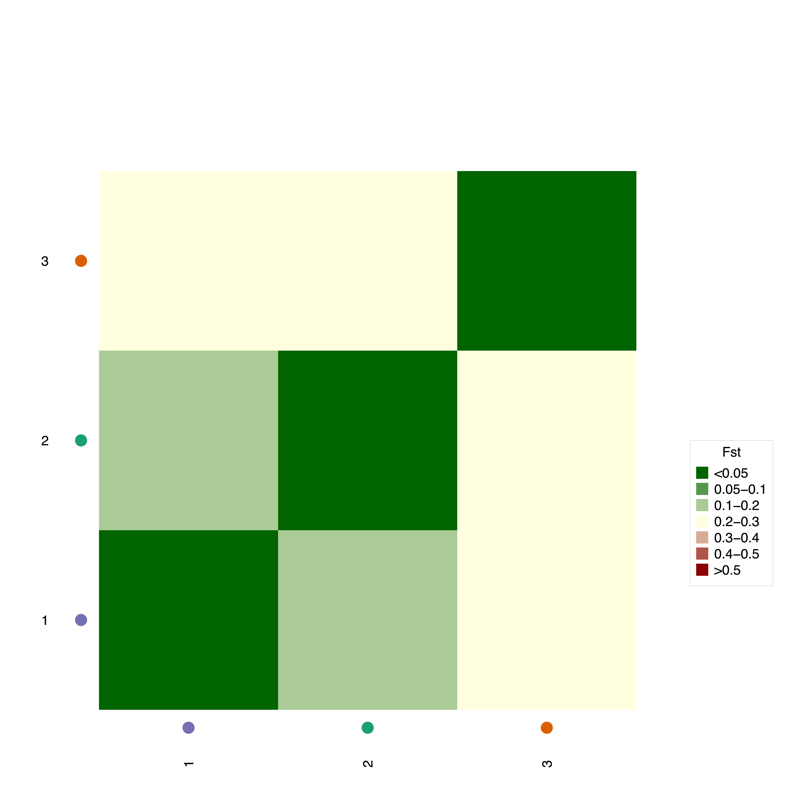 |
| **iii)**  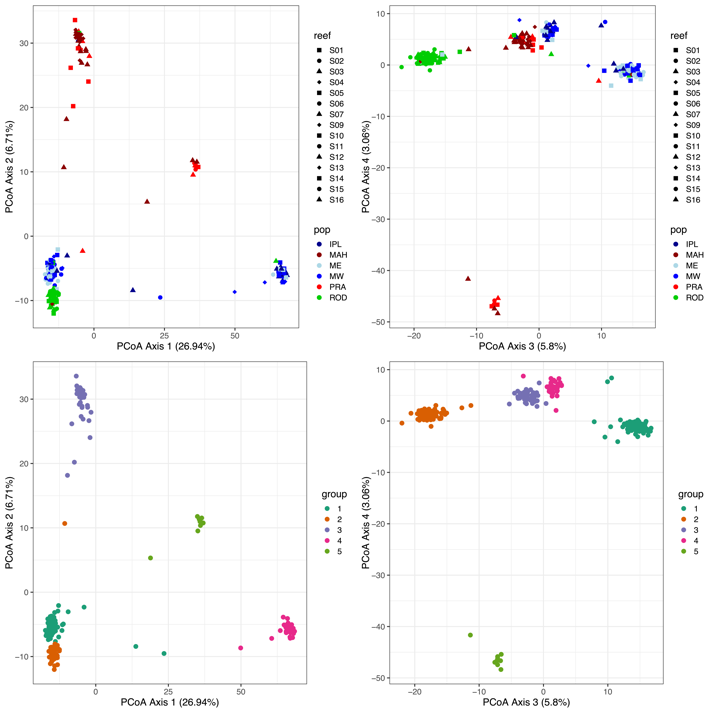 | **iv)**  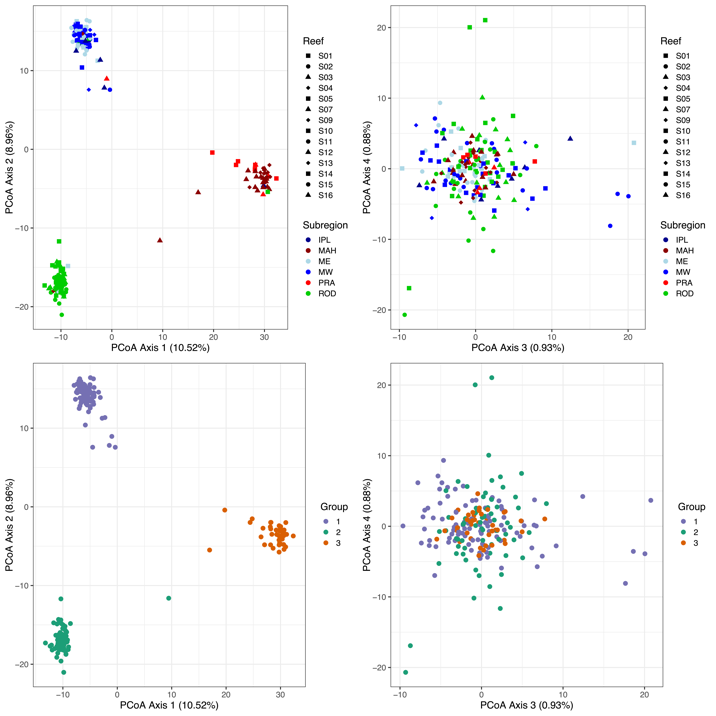 | |

**(b) *Pocillopora damicornis***

| **Pre-filtering** | **Post-filtering** | |
| --- | --- | --- |
| **i)**  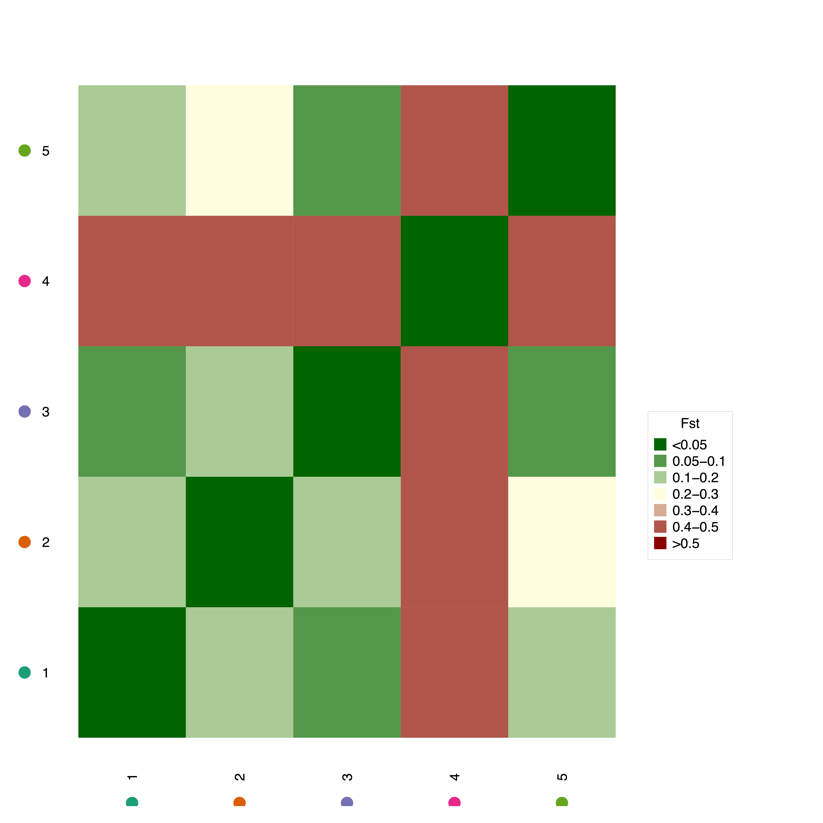 | | **ii)**  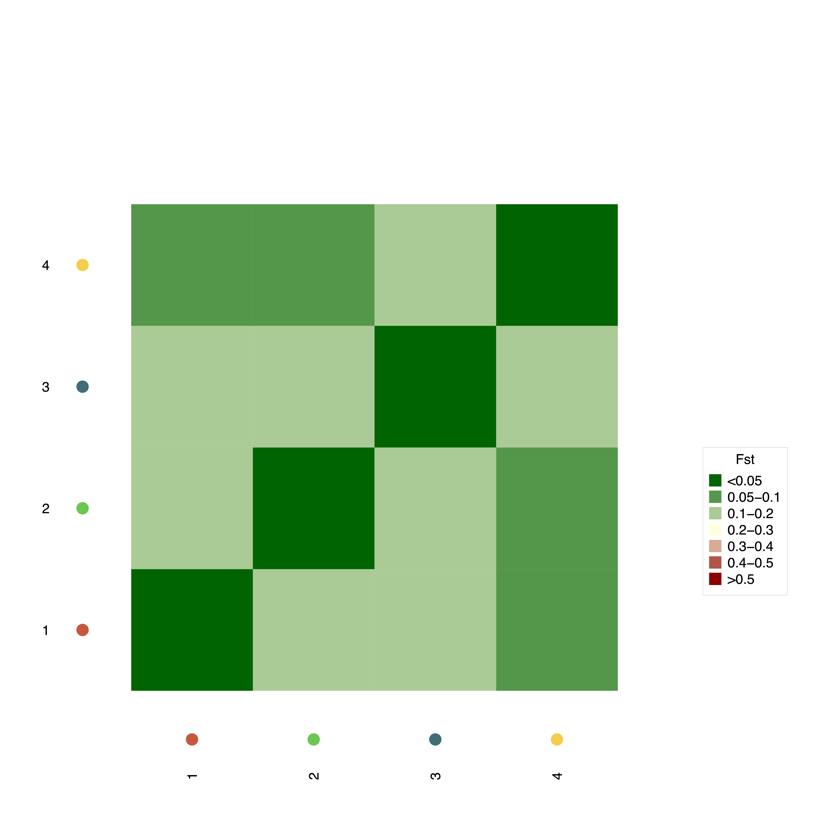 |
| **iii)**  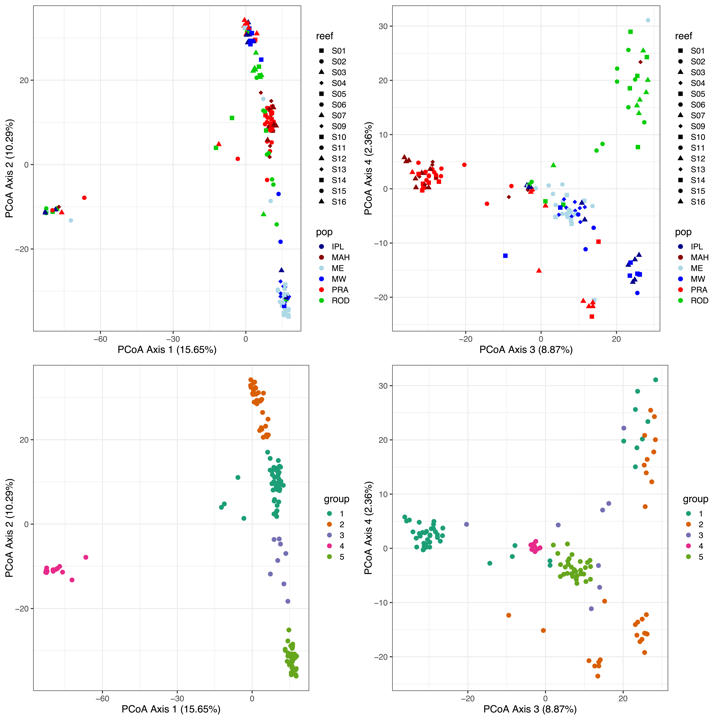 | **iv)**  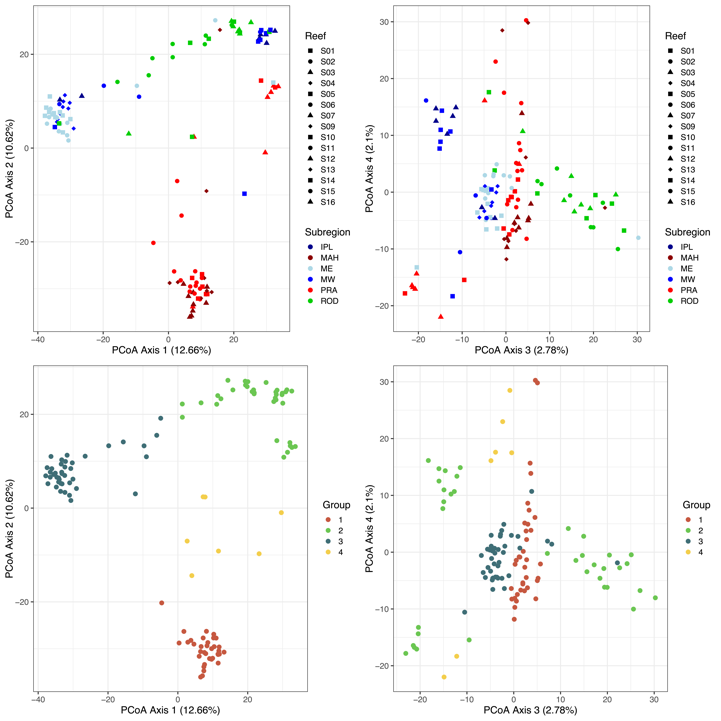 | |

**Suppl. Fig. S4.** **AMOVA histograms.** Analysis of molecular variance (AMOVA) to detect population differentiation using SNP markers comparing allele frequency variation of SNPs for (**a**) *Acropora muricata* and (**b**) *Pocillopora damicornis*. Histograms represent the distribution of randomly permutated values (grey boxes) compared to the observed results (black line) for SNP allele frequency variation within samples, between samples, between sample sites, between subregions, and between regions.

**(a)**

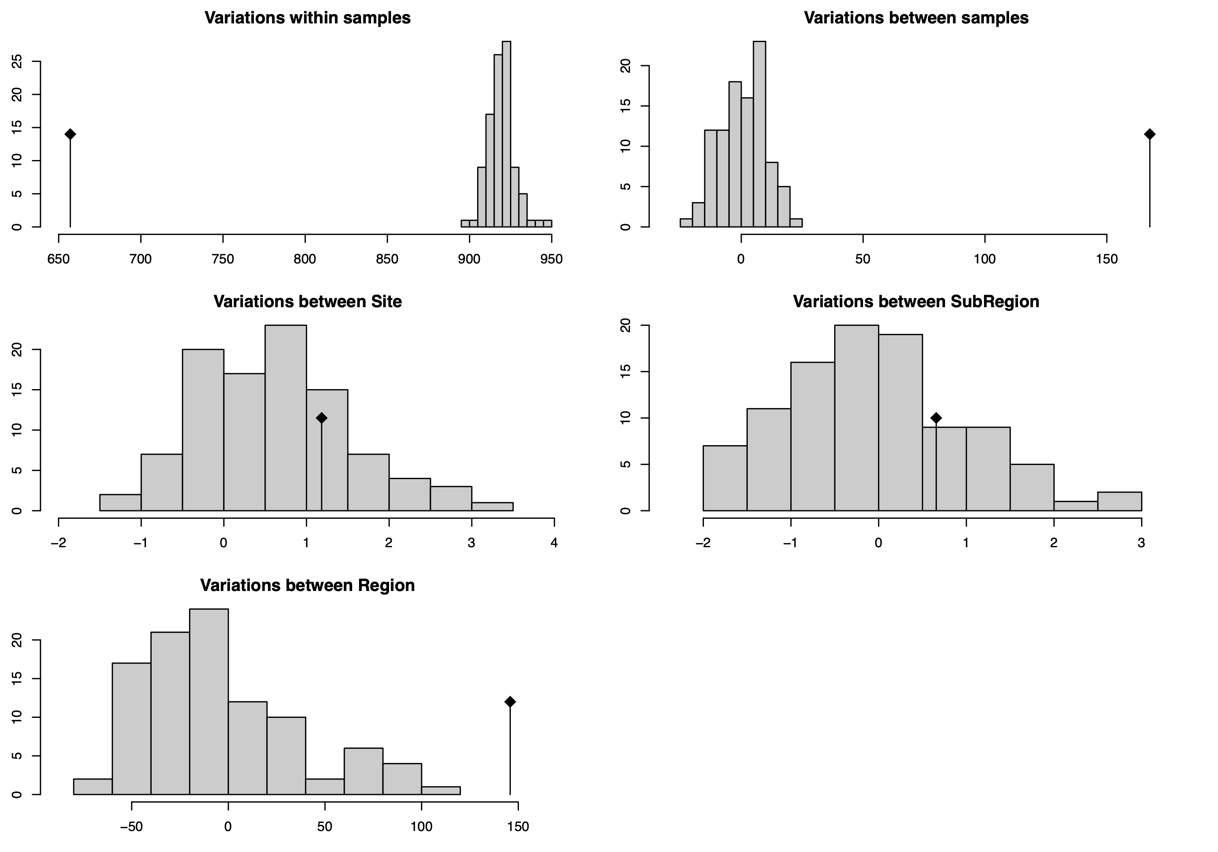

**(b)**

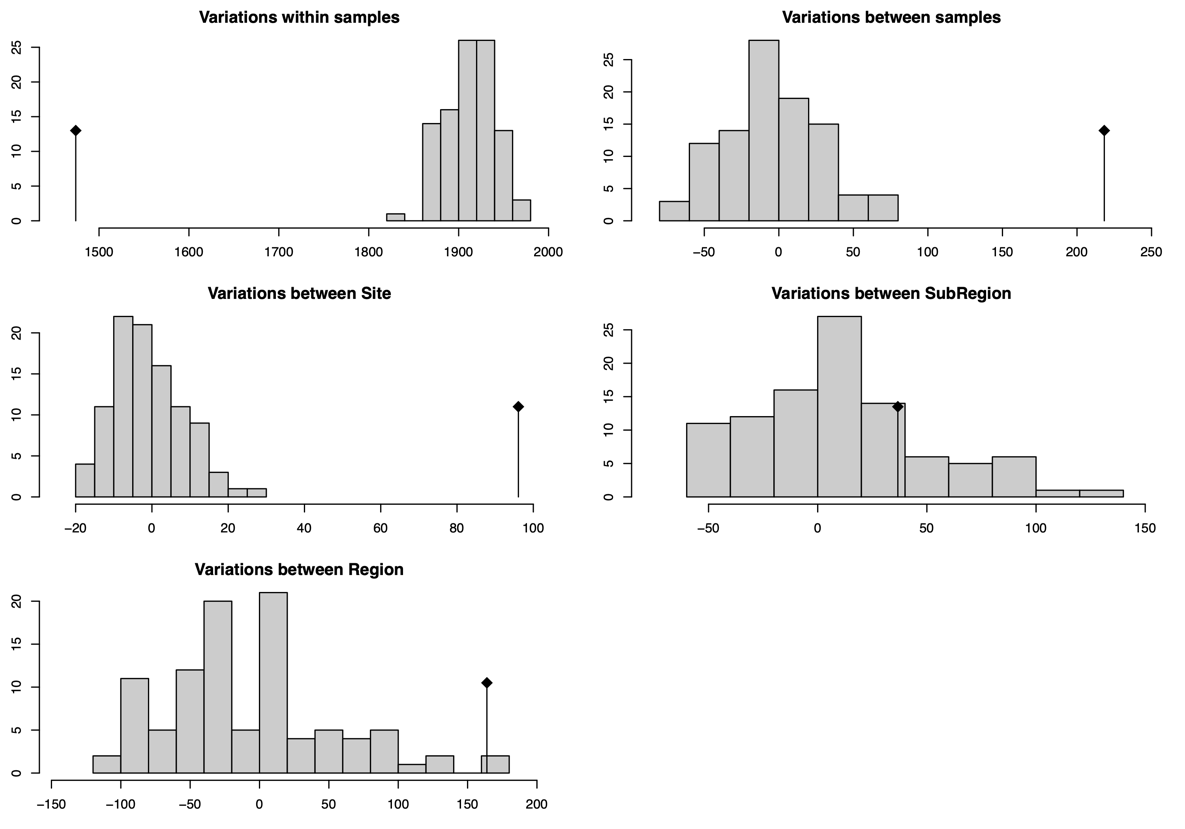

**Suppl. Fig. S5.** **Adaptive and connectivity indices.** Spatial representations of adaptive potentials and connectivity metrics for **(a)** *A. muricata* and **(b)** *P. damicornis* at coral reefs across the West Indian Ocean. **Top left:** The Adaptive Potential Index (API) indicating the area (km^2^) of reefs upstream (ICI) with heat adapted corals (Adaptive Score >0.8). **Top right:** The Adaptive Score (AS) indicating the relative level of adaptation to thermal stressors (sSST and mDHW) from 0-1 (low to high adaptive values). **Bottom left:** Outbound Connectivity Index (OCI) indicating the area (in km^2^) of reef downstream of the target reef a sea distance corresponding to genetic connectivity of F_ST_<0.1. **Bottom right**: Inbound Connectivity Index (ICI) indicating the area (in km^2^) of reef upstream of the target reef a sea distance corresponding to genetic connectivity of F_ST_<0.1.

**(a)**

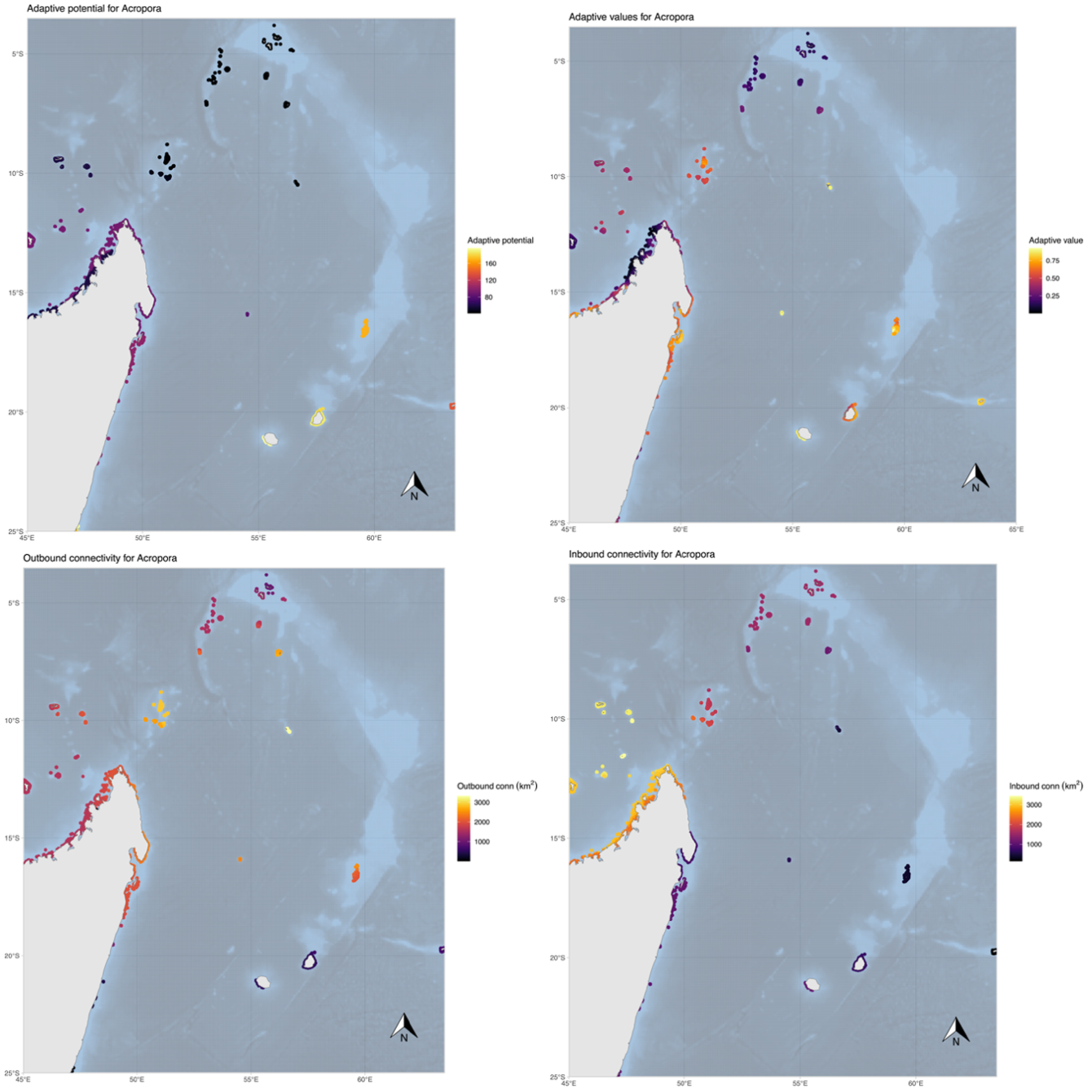

**(b)**

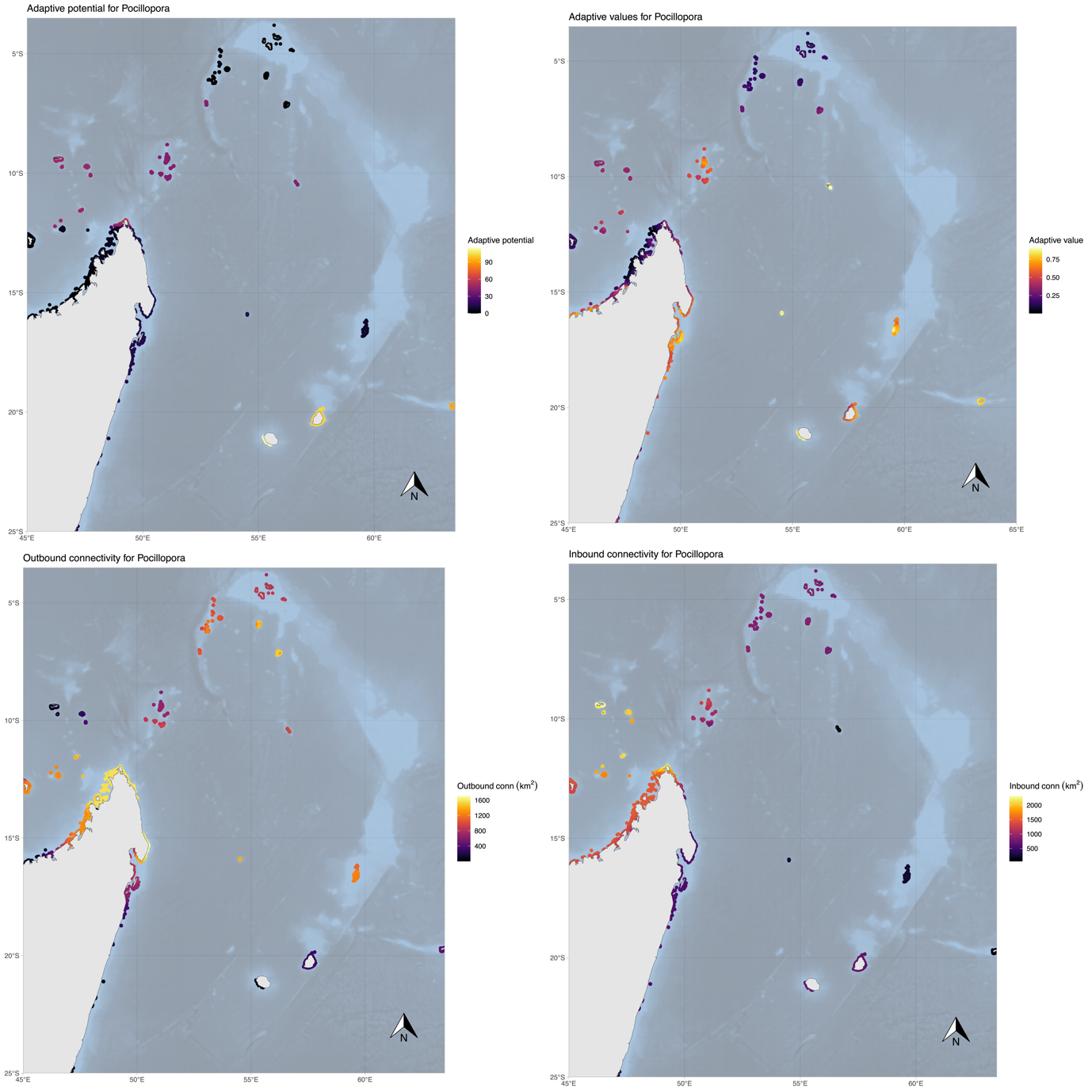

**Suppl. Table S1.** **Sample sites and sample sizes.** Summary of the 15 sample sites where *Acropora muricata* and *Pocillopora damicornis* colonies were collected for genotyping in the West Indian Ocean. Summarised in the table: Region, Sub region, reef name, average latitude (LAT) and longitude (LON), diving depth in meters (Depth), and the number of colonies remaining after SNP genotype filtering.

Region abbreviations: MAU = Maurice, ROD = Rodrigues, SEY = Seychelles. Sub-region abbreviations: IPL = Ile Plate, ME = East Mauritius, MW = West Mauritius, ROD = Rodrigues, MAH = Mahé, PRA = Praslin.

| **Code** | **Region** | **Sub-region** | **Reef name** | **LAT** | **LON** | **Depth (m)** | **Number of *A. muricata*** | | **Number of *P. damicornis*** | |
| --- | --- | --- | --- | --- | --- | --- | --- | --- | --- | --- |
|  |  |  |  |  |  |  | Sampled | Post-filtering | Sampled | Post-filtering |
| S12 | MAU | IPL | Ile Plate | -19.881 | 57.668 | 1-3 | 29 | 8 | 29 | 7 |
| S02 | MAU | ME | Belle Mare | -20.192 | 57.781 | 1-3 | 29 | 25 | 29 | 12 |
| S10 | MAU | ME | Gd Bourg Melville | -20.010 | 57.687 | 1-3 | 29 | 13 | 29 | 11 |
| S11 | MAU | MW | Balaclava | -20.077 | 57.510 | 1-3 | 29 | 23 | 28 | 4 |
| S13 | MAU | MW | FlicFlac | -20.277 | 57.365 | 1-3 | 29 | 5 | 29 | 9 |
| S01 | MAU | MW | St Felix | -20.512 | 57.465 | 1-3 | 29 | 19 | 29 | 7 |
| S16 | ROD | ROD | 80 Brisants | -19.833 | 63.342 | NA | 26 | 17 | 28 | 7 |
| S15 | ROD | ROD | Gd Bassin Bretagne | -19.659 | 63.340 | NA | 29 | 24 | 27 | 8 |
| S14 | ROD | ROD | SEMPA | -19.739 | 63.479 | NA | 28 | 27 | 27 | 7 |
| S09 | SEY | MAH | Anse La Mouche | -4.739 | 55.474 | NA | 13 | 10 | 21 | 3 |
| S04 | SEY | MAH | Baie Ternay | -4.640 | 55.377 | 1-3 | 8 | 5 | 29 | 6 |
| S03 | SEY | MAH | Ile Au Cerf | -4.635 | 55.491 | 1-3 | 29 | 19 | 29 | 9 |
| S07 | SEY | PRA | Coco | -4.369 | 55.853 | 3-10 | 13 | 4 | 13 | 7 |
| S05 | SEY | PRA | Cousine | -4.351 | 55.651 | 3-10 | 12 | 6 | 28 | 9 |
| S06 | SEY | PRA | Curieuse | -4.284 | 55.740 | 3-10 | 13 | 0 | 28 | 14 |
|  |  |  |  |  |  | **Total** | **345** | **205** | **403** | **120** |

**Suppl. Table S2.** **Environmental variable description.** Variables used to characterise the reefs around the study regions of Mauritius, Rodrigues and the Seychelles in the West Indian Ocean. Provided are: acronym used in the main text, full variable name, calculated spatial resolution, calculated temporal window, open-access website from which variables are sourced (RECIFS: Selmoni et al., 2020, or the Allen Coral Atlas (ACA): Allen Coral Atlas 2022), details of the variable, and the origin of the raw data used to make the variables.

| **Acronym** | **Variable** | **Spatial resolution** | **Temporal window** | **Repository** | **Details** | **Source** |
| --- | --- | --- | --- | --- | --- | --- |
| DEP | Depth | 5x5km | n/a | RECIFS | Average depth around reef calculated with a 5km buffer | Global Multi-Resolution Topography Data Synthesis |
| mDHW | Degree Heating Week (mean) | 5x5km | Monthly max from 1985-2021 | RECIFS | Overall mean of the monthly maxima DHW (accumulated thermal stress over previous 12 weeks | NOAA Coral Reef Watch |
| sDHW | Degree Heating Week (sd) | 5x5km | Monthly max from 1985-2021 | RECIFS | Overall sd of the monthly maxima DHW (accumulated thermal stress over previous 12 weeks | NOAA Coral Reef Watch |
| mSST | Sea Surface Temperature (mean) | 5x5km | Monthly ave from 1985-2021 | RECIFS | Overall mean of the monthly SST | NOAA Coral Reef Watch |
| sSST | Sea Surface Temperature (sd) | 5x5km | Monthly ave from 1985-2021 | RECIFS | Overall sd of the monthly SST | NOAA Coral Reef Watch |
| mCHL | Chlorophyll concentration (mean) | 5x5km | Monthly ave from 1997-2021 | RECIFS | Overall mean of the monthly average of mass chlorophyll-a concentration in seawater | Copernicus Marine Service |
| sCHL | Chlorophyll concentration (sd) | 5x5km | Monthly ave from 1997-2021 | RECIFS | Overall sd of the monthly average of mass chlorophyll-a concentration in seawater | Copernicus Marine Service |
| mSPM | Suspended Particulate Matter (mean) | 5x5km | Monthly ave from 1997-2021 | RECIFS | Overall mean of the mass concentration of suspended matter in seawater | Copernicus Marine Service |
| sSPM | Suspended Particulate Matter (sd) | 5x5km | Monthly ave from 1997-2021 | RECIFS | Overall sd of the mass concentration of suspended matter in seawater | Copernicus Marine Service |
| **Acronym** | **Variable** | **Spatial Res** | **Temporal window** | **Source** | **Details** | **Origin of raw data** |
| CROP | Density of cropland | 5x5km | Measured in 2015–2019 | RECIFS | Percentage of pixels in 5km buffer that correspond to cropland cover | Copernicus Global Land Service |
| LAND | Density of land surface | 5x5km | n/a | RECIFS | Percentage of pixels in 5km buffer that correspond to land cover | Global Multi-Resoulution Topography Data Synthesis |
| HPOP | Human population along coastline | 5x5km | Measured in 2000, 2005, 2010, 2015, 2020 | RECIFS | Mean human population density in the 5km buffer around the reef of interest | Center for International Earth Science Information |
| BOAT | Boat detection | 5x5km | Measured in 2017–2021 | RECIFS | Mean of the percentage of boats detected using satellite imagery | Earth Observation Group, Payne Institute for Public Policy |
| ReefSlope | Prop. reef slope 250m around reef | 10m | n/a | ACA | Submerged, sloping area extending seaward from the Reef Crest (or Flat) towards the shelf break. | Planet Dove satellite imagery |
| ReefFlat | Prop. reef flat 250m around reef | 10m | n/a | ACA | Adjacent to the seaward edge of the reef, Outer Reef Flat is a levelled (near horizontal) broad and shallow carbonate platform, displaying distinct wave-driven zonation. | Planet Dove satellite imagery |
| BackReef | Prop. back reef 250m around reef | 10m | n/a | ACA | A complex, interior - often gently sloping - reef zone occurring behind the Reef Flat. Of variable depth (but deeper than Reef Flat and more sloped), it is sheltered, sediment-dominated and often punctuated by coral outcrops. | Planet Dove satellite imagery |
| Plateau | Prop plateau 250m around reef | 10m | n/a | ACA | Deep submerged (> 5 m), hard-bottomed, horizontal to gently sloping (angle shallower than 10 ° approx), seaward facing reef platform. | Planet Dove satellite imagery |

**Suppl. Table S3. Species ID** **Genetic confirmation.** Genetic confirmation of retained *Pocillopora* samples (i.e., after filtering for clones and cryptic individuals) as *P. damicornis,* using DArT sequences within the 18S and ITS-2 ribosomal DNA (rDNA) regions. Five individuals from each of the four *Pocillopora* PCoA genetic groups (Suppl. Fig. S3B) were randomly selected for analysis. These 20 individuals were queried against rDNA sequences obtained from NCBI (Genbank ID) for four target *Pocillopora* species (*P. damicornis*, P*. verrucosa, P. eydouxi*, *P. elegans*) and two outgroup species (*Stylophora pistillata* and *Seriatopora* sp.). The Percentage of Identity indicates how much our 20 samples aligned to the target species for each queried sequence, alongside the length of alignment (base pairs; bp) and the expectation value (E-value) of the BLAST hits. As the sequences for all 20 individuals at these loci were 100% in alignment with each other, here we show results from the genotyped individual 3586603 (ID ‘01_P_12’ from Site S01 in Mauritius). Nucleotide alignments for the 18S and ITS-2 sequences of the 20 individuals alongside three cryptic individuals and target species are shown below (aligned using ClustalW).

| **DNA region** | **Query sequence** | **Target sequence (Genbank)** | **Target species** | **Percentage of identity** | **Alignment length (Bp)** | **E-value** |
| --- | --- | --- | --- | --- | --- | --- |
| 18S | 3586603_18S | PQ434668.1 | *Pocillipora damicornis* | 100.00 | 105 | 1.42e^-54^ |
| 18S | 3586603_18S | XR_010716799.1 | *Pocillopora verrucosa* | 98.53 | 67 | 8.76e^-32^ |
| 18S | 3586603_18S | LT631145.1 | *Stylophora pistillata* | 95.24 | 105 | 3.09e^-46^ |
| 18S | 3586603_18S | LT631140.1 | *Seriatopora* sp. | 95.24 | 105 | 3.09e^-46^ |
| ITS-2 | 3586603_ITS2 | KF846519.1 | *Pocillopora damicornis* | 99.06 | 108 | 5.25e^-54^ |
| ITS-2 | 3586603_ITS2 | HM013854.1 | *Pocillopora eydouxi* | 96.26 | 108 | 5.29e^-49^ |
| ITS-2 | 3586603_ITS2 | EU314802.1 | *Pocillopora elegans* | 96.26 | 108 | 5.29e^-49^ |
| ITS-2 | 3586603_ITS2 | OK448746.1 | *Pocillopora verrucosa* | 94.29 | 36 | 2.59e^-12^ |

18S rDNA Alignment

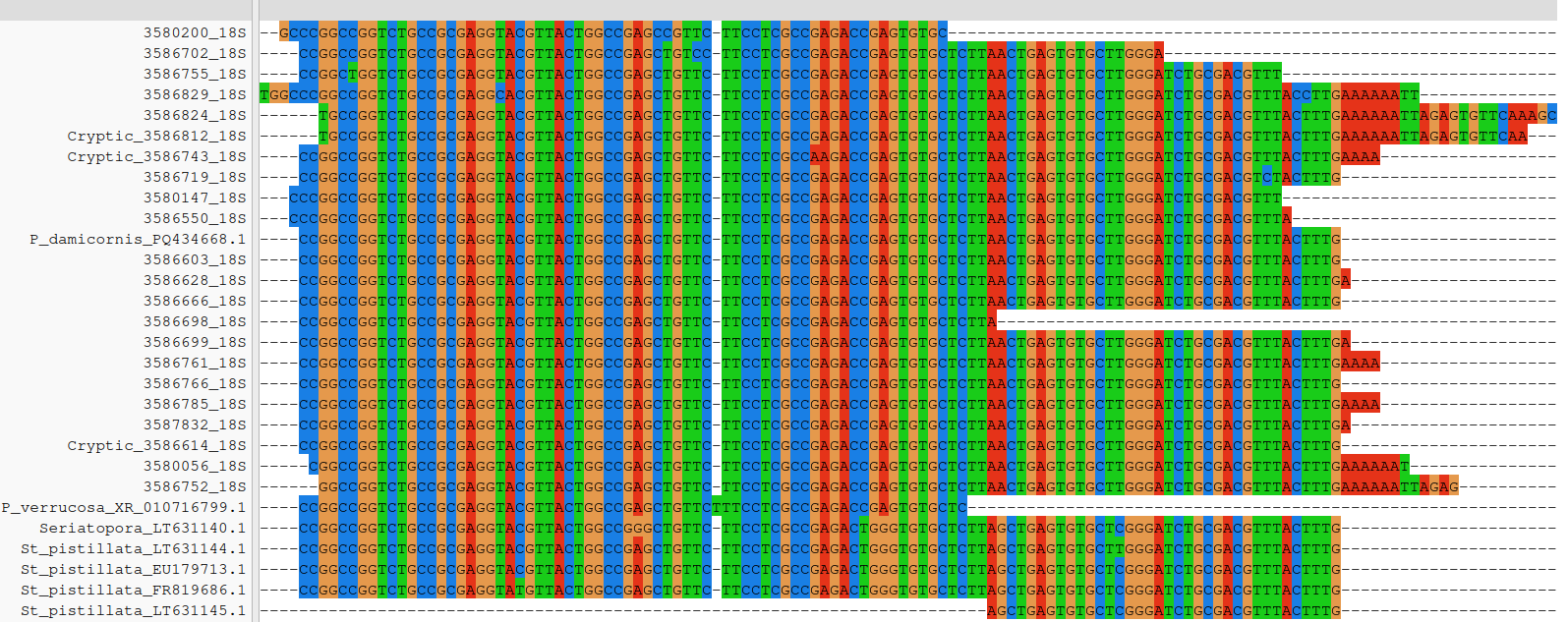

ITS-2 rDNA alignment

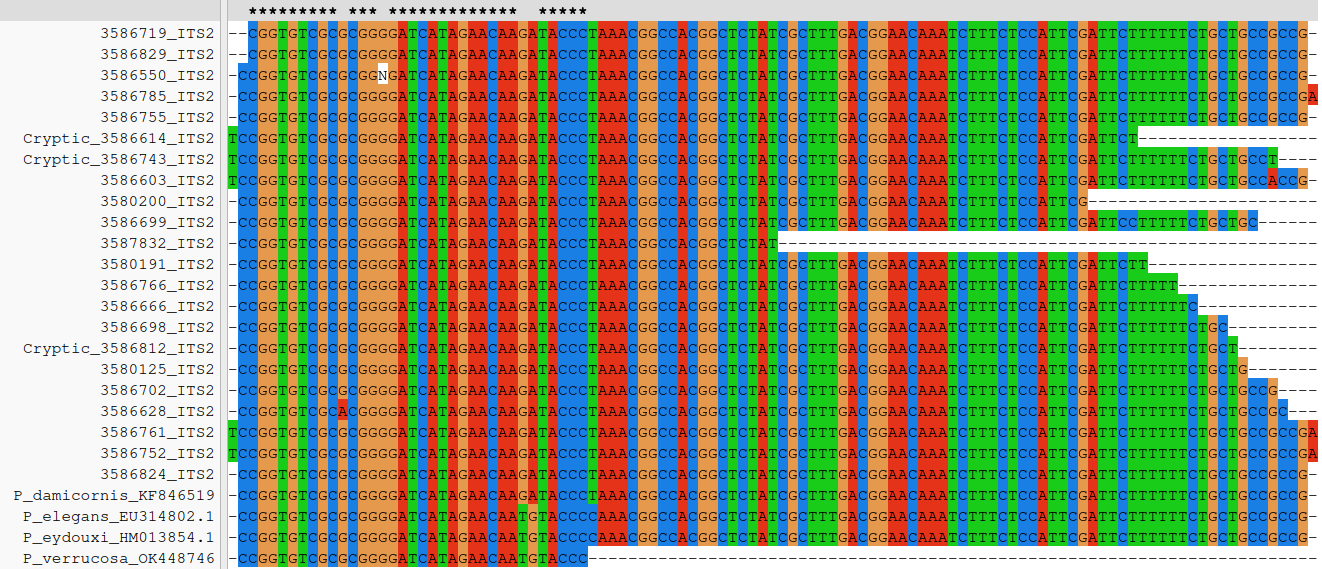

**Suppl. Table S4.** **Genomic filtering.** Number of SNPs and individuals retained at each step of genomic filtering for *Acropora muricata* and *Pocillopora damicornis* at all sampled reefs of the WIO. The first row indicates the number of SNPs and individuals available from DArT-seq genotyping, followed by numbers retained after filtering for clones (relatedness >0.95), missingness of SNPs (MNsnp; <0.8), missingness for individuals (MNind; <0.8), minor allele frequency (MAF; >0.05) and cryptic individuals. Filtering for missingness and MAF were then performed again following the removal of cryptic individuals (MNsnp 2, MNind 2, MAF 2). We also show the SNPs remaining had we undertaken LD pruning (shaded out row), where we decided to retain all SNPs for the downstream analyses. Finally, we show the number of SNPs detected as Outlier Loci using *PCAdapt* and the number of SNPs retained for the neutral dataset.

| **Filtering step** | ***Acropora muricata*** | | ***Pocillopora damicornis*** | |
| --- | --- | --- | --- | --- |
|  | **SNPs** | **Individuals** | **SNPs** | **Individuals** |
| **DArT-seq** | 73,253 | 345 | 65,708 | 403 |
| **Blast and Clone removal** | 63,511 | 297 | 52,219 | 142 |
| **MNsnp 1** | 15,007 | 297 | 26,764 | 142 |
| **MNind 1** | 14,282 | 248 | 26,754 | 134 |
| **MAF 1** | 8,043 | 248 | 14,755 | 134 |
| **Cryptic individuals** | 8,043 | 205 | 14,755 | 120 |
| **MNsnp 2** | 7,772 | 205 | 13,903 | 120 |
| **MNind 2** | 7,772 | 205 | 13,903 | 120 |
| **MAF 2** | 5,757 | 205 | 12,953 | 120 |
| **LD pruning** | 3,309 | 205 | 7,171 | 120 |
| **Outlier loci** | 74 | 205 | 138 | 120 |
| **Neutral loci** | 5,683 | 205 | 12,815 | 120 |

**Suppl. Table S5.** **Population statistics.** Population statistics for (**a**) *Acropora muricata* and (**b**) *Pocillopora damicornis* based on neutral SNP genotype matrices. Statistics are calculated per sampled reef, where we report observed heterozygosity (Ho), expected heterozygosity (He), unbiased expected heterozygosity (uHe; corrected by *n* individuals – nInd), and the inbreeding coefficient (F_IS_; values >0.2 are highlighted in yellow).

**(a)**

| **Region** | **Sub Region** | **Reef** | **nInd** | **Ho** | **He** | **uHe** | **F_IS_** |
| --- | --- | --- | --- | --- | --- | --- | --- |
| **Seychelles** | PRA | S05 | 5.5 | 0.25 | 0.26 | 0.29 | 0.14 |
|  |  | S07 | 3.5 | 0.19 | 0.24 | 0.28 | 0.33 |
|  | MAH | S03 | 17.5 | 0.22 | 0.28 | 0.29 | 0.23 |
|  |  | S04 | 4.7 | 0.20 | 0.24 | 0.26 | 0.24 |
|  |  | S09 | 9.3 | 0.21 | 0.29 | 0.30 | 0.32 |
| **Mauritius** | IPL | S12 | 7.5 | 0.25 | 0.29 | 0.31 | 0.20 |
|  | MW | S01 | 18.3 | 0.22 | 0.28 | 0.28 | 0.21 |
|  |  | S11 | 22.2 | 0.25 | 0.29 | 0.29 | 0.13 |
|  |  | S13 | 4.7 | 0.24 | 0.26 | 0.29 | 0.18 |
|  | ME | S02 | 24.2 | 0.24 | 0.28 | 0.29 | 0.18 |
|  |  | S10 | 12.5 | 0.24 | 0.28 | 0.29 | 0.17 |
| **Rodrigues** | ROD | S14 | 25.8 | 0.23 | 0.28 | 0.29 | 0.22 |
|  |  | S15 | 22.9 | 0.22 | 0.27 | 0.28 | 0.22 |
|  |  | S16 | 16.3 | 0.22 | 0.27 | 0.28 | 0.23 |

**(b)**

| **Region** | **Sub Region** | **Reef** | **nInd** | **Ho** | **He** | **uHe** | **F_IS_** |
| --- | --- | --- | --- | --- | --- | --- | --- |
| **Seychelles** | PRA | S06 | 13.5 | 0.23 | 0.26 | 0.27 | 0.13 |
|  |  | S05 | 8.7 | 0.22 | 0.25 | 0.27 | 0.19 |
|  |  | S07 | 6.5 | 0.17 | 0.23 | 0.25 | 0.30 |
|  | MAH | S03 | 8.7 | 0.22 | 0.23 | 0.24 | 0.07 |
|  |  | S04 | 5.8 | 0.22 | 0.24 | 0.27 | 0.16 |
|  |  | S09 | 2.9 | 0.29 | 0.22 | 0.27 | -0.07 |
| **Mauritius** | IPL | S12 | 6.7 | 0.20 | 0.24 | 0.26 | 0.23 |
|  | MW | S01 | 6.8 | 0.21 | 0.24 | 0.26 | 0.20 |
|  |  | S11 | 3.9 | 0.27 | 0.26 | 0.30 | 0.08 |
|  |  | S13 | 8.7 | 0.24 | 0.23 | 0.25 | 0.03 |
|  | ME | S02 | 11.5 | 0.23 | 0.25 | 0.26 | 0.12 |
|  |  | S10 | 10.6 | 0.22 | 0.24 | 0.26 | 0.12 |
| **Rodrigues** | ROD | S14 | 6.7 | 0.24 | 0.26 | 0.28 | 0.15 |
|  |  | S15 | 7.8 | 0.26 | 0.25 | 0.27 | 0.05 |
|  |  | S16 | 6.7 | 0.24 | 0.24 | 0.26 | 0.07 |
