## Supplementary material for "Coral genetic structure in the Western Indian Ocean mirrors ocean circulation and thermal stress": Suppl. Methods: Selection of sample sites

**Supplementary material:** Selection of sample sites using multivariate environmental characterization

This supplementary material summarizes the baseline assessment of the environmental conditions in the reefs of the region to establish a sampling strategy for seascape genomics analyses.

### ****Sampling design in seascape genomics****

The goal of a seascape genomics study is to uncover genetic variants in a given population that are associated with environmental gradients of the seascape where the population lives (Riginos et al., 2016; Selmoni et al., 2020a). In the marine environment, and particularly in proximity of the coastline, environmental gradients are often superposed to each other, and this is an aspect that hampers the detection of adaptive signals. To overcome this issue, it is crucial to design a sampling strategy that minimizes collinearity between environmental variables.

Here, we follow the approach of Selmoni et al. (2020) to optimize the selection of sample sites across the regions of interest, which has been successfully used in coral seascape genomic analyses to reveal candidate loci under thermal selection in New Caledonia (Selmoni et al., 2021).

This approach is based on a classification of the possible sampling locations through a multivariate environmental characterization. The final goal of this approach is to split the reefs of a study area in different environmental zones showing distinct environmental conditions. Sampling sites can then be established in different environmental zones to maximize the environmental variability captured by the seascape genomics study.

### Multivariate environmental characterization

To elaborate a solid sampling strategy, it is crucial to have a meaningful environmental description of the reefs of the different study areas. We performed this task by using datasets derived from remote sensing (Table 1) since they allow us to (1) perform a standardized comparison between distant sites/islands; (2) include a large number of different environmental variables (3) compute climatologies over several decades, a timeframe that is relevant to detect adaptive processes.

It is important to point out that one of the main limitations of remote sensing datasets is the fact that they overlook fine-scale environmental variation (i.e. at the scale of a few hundreds of meters). To mitigate this issue, we used environmental datasets at the lowest resolution available for every variable (~ 5 km). For some of the variables, the spatial resolution is clearly non-optimal to detect adaptive signals within a single island/region (e.g. for pH spatial resolution is 25km). However, such variables can still be useful to uncover adaptive processes occurring across different islands/regions.

Two main types of variables were retrieved from publicly available databases: those describing changes in the water conditions across time/seasons (e.g. sea surface temperature, chlorophyll concentration, water velocity, etc.); and those that describe the physical structure of the seascape (e.g. bathymetry) or the land-use along the coastline (e.g. population density, boat detection). For the first type of variables, we computed two overall statistics: overall average and overall standard deviation. For the other variables, we computed metrics characterizing the seascape structure (e.g. average bathymetry) or the land-use (e.g. population density) in a radius of 10 and 50 km around a reef of interest.

| **Table 1** Environmental variables used to describe seascape for selecting study sites***.*** | | | |
| --- | --- | --- | --- |
| **Environmental Condition** | **Temporal Range** | **Resolution** | **Source** |
| Sea Surface Temperature anomalies, Degree heating week | 1985-2020, daily | 5 km | Coral Reef Watch (NOAA) |
| Salinity | 2007-2020, daily | 8 km | COPERNICUS |
| Water Currents | 2007-2020, daily | 8 km | COPERNICUS |
| Iron concentration | 1993-2020 | 25 km | COPERNICUS |
| Oxygen concentration | 1993-2020 | 25 km | COPERNICUS |
| pH | 1993-2020 | 25 km | COPERNICUS |
| Phosphate concentration | 1993-2020 | 25 km | COPERNICUS |
| Chlorophyll Concentration | 1997-2020, monthly | 4 km | COPERNICUS |
| Suspended Particulate Matter | 1997-2020, monthly | 4 km | COPERNICUS |
| Landcover (to compute proximity to urban areas) | 2020 | 1 km | COPERNICUS  (Global Land Services) |
| Population Density | 2020 | 1 km | NASA |
| Bathymetry | - | 1 km-100m | MGDS |
| Boat detection | 2020 | 1 km | NOAA (Earth Observation Group) |

### Defining environmental zones

For each of the four regions of interest (Maurititus, Mahé (SEY), Praslin (SEY) and Rodrigues), the definition of the environmental zones was performed as follows. First, we accessed the UNEP inventory the reefs of the world (<https://data.unep-wcmc.org/datasets/1>) and retrieved the position of all the reefs from the regions of interest. Next, we reported the corresponding reef-area into a regular grid of thousands of cells of size 500x500m. For every reef-cell, we then retrieved the respective values from the multivariate environmental characterization. Next, we performed a Principal Component Analysis (PCA) to summarize the environmental variation across the different reef-cells. The PCA was performed separately for three types of variables:

1) **Heat:** such as averages and variations of SST, and variables describing accumulation of heat such as Degree-heating-week.

2) **Water chemistry:** including averages and variations of water pH, salinity, turbidity, oxygen concentration, phosphate concentration.

3) **Seascape configuration**: these are variables that define the surrounding of a reef, for example the frequency of surface land, the average depth, the proximity to urban areas.

Based on the results of the three PCAs, we performed a hierarchical clustering separating the different reef-cell according to their environmental characteristics. The number of K clusters computed reflected the number of target sites required as determined by stakeholders (K=5 for Mauritius; K=3 for Rodrigues; K=4 for Mahé; K=4 for Praslin).

### Selecting sample sites based on environmental zones

Maps of environmental zones were created for each region of interest, presented at the end of this document. Below each map are tables that show the rates of environmental variation between the different zones for some of the variables used in the multivariate environmental characterization.

For each region, one sampling location was selected within each environmental zone, resulting in 15 reefs (5 in Mauritius; 3 in Rodrigues; 4 in Mahé, SEY; 4 in Praslin, SEY). The exact location was decided in collaboration with local stakeholders of the regions, following criteria including: facility of access, presence of study species, depth profile (to be consistent between all the sampling locations; max. 10m depth), relevance for conservation plans (e.g. nursery, protected area) and for socio-economic activities.

**Mauritius**

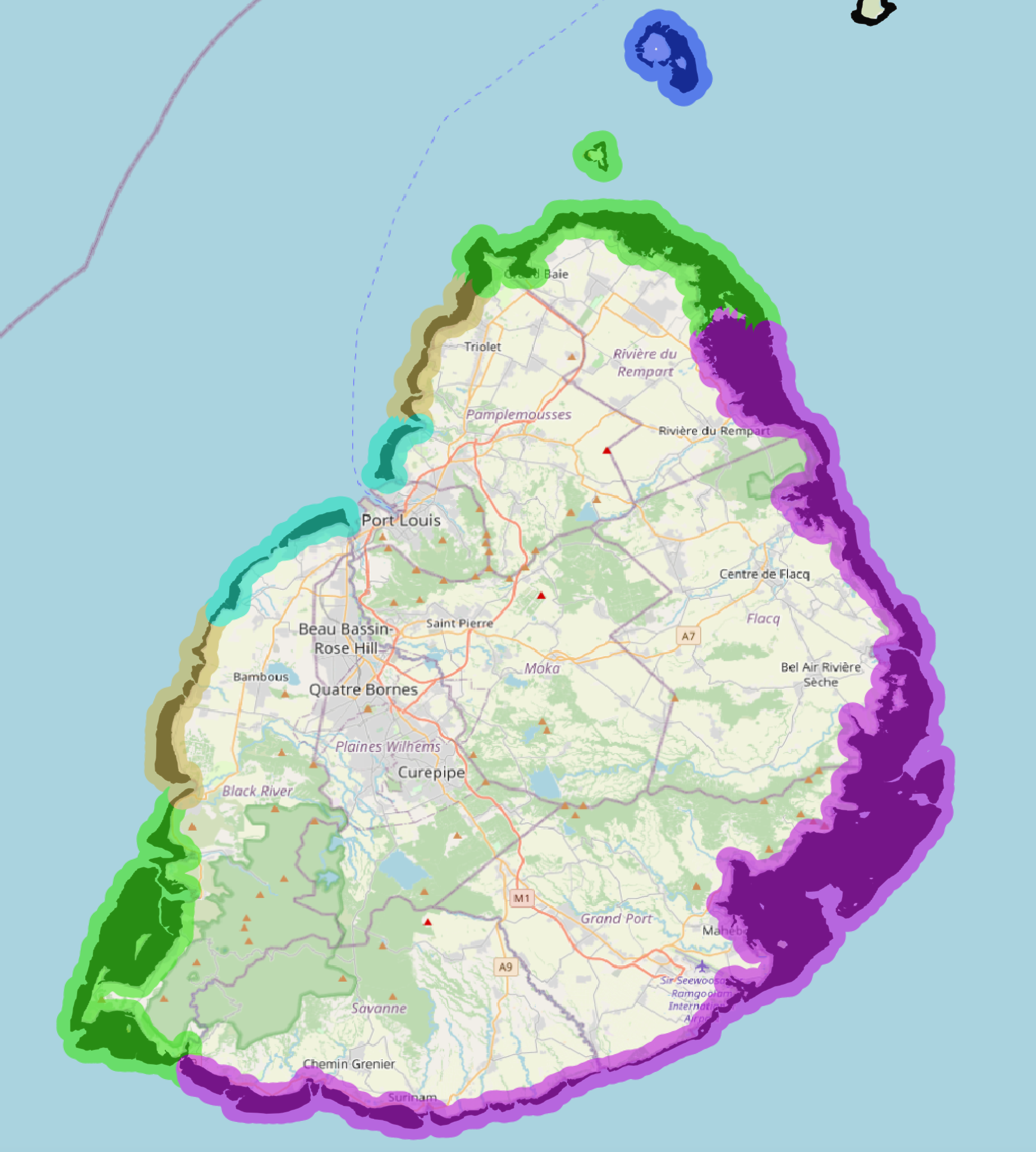

| ENV_zone | Degree heating week | SST st deviation | Chlorophyll mean | pH mean | Sea current velocity st. deviation | Depth in a 10 km buffer | Boat detection in a 10 km buffer | Population density in a 10 km buffer |
| --- | --- | --- | --- | --- | --- | --- | --- | --- |
| 1 | 0.454±0.046 | 1.74±0.015 | 0.238±0.065 | 8.07±0.00026 | 0.0773±0.013 | -355±220 | 0.00331±6e-04 | 227±190 |
| 2 | 0.375±0.026 | 1.73±0.011 | 0.145±0.032 | 8.07±4.3e-05 | 0.0634±0.0099 | -952±200 | 0.0212±0.025 | 528±290 |
| 3 | 0.587±0.044 | 1.74±0.0055 | 0.336±0.2 | 8.07±5e-04 | 0.0784±0.017 | -386±290 | 0.00404±0.0031 | 323±100 |
| 4 | 0.357±0.016 | 1.73±0.004 | 0.249±0.12 | 8.07±2.5e-05 | 0.0637±0.0052 | -957±190 | 0.0874±0.011 | 1150±240 |
| 5 | 0.454±0.0077 | 1.72±0.00055 | 0.101±0.0068 | 8.07±3.9e-05 | 0.0877±0.0087 | -168±50 | 0±0 | 6.46±13 |

**Rodrigues**

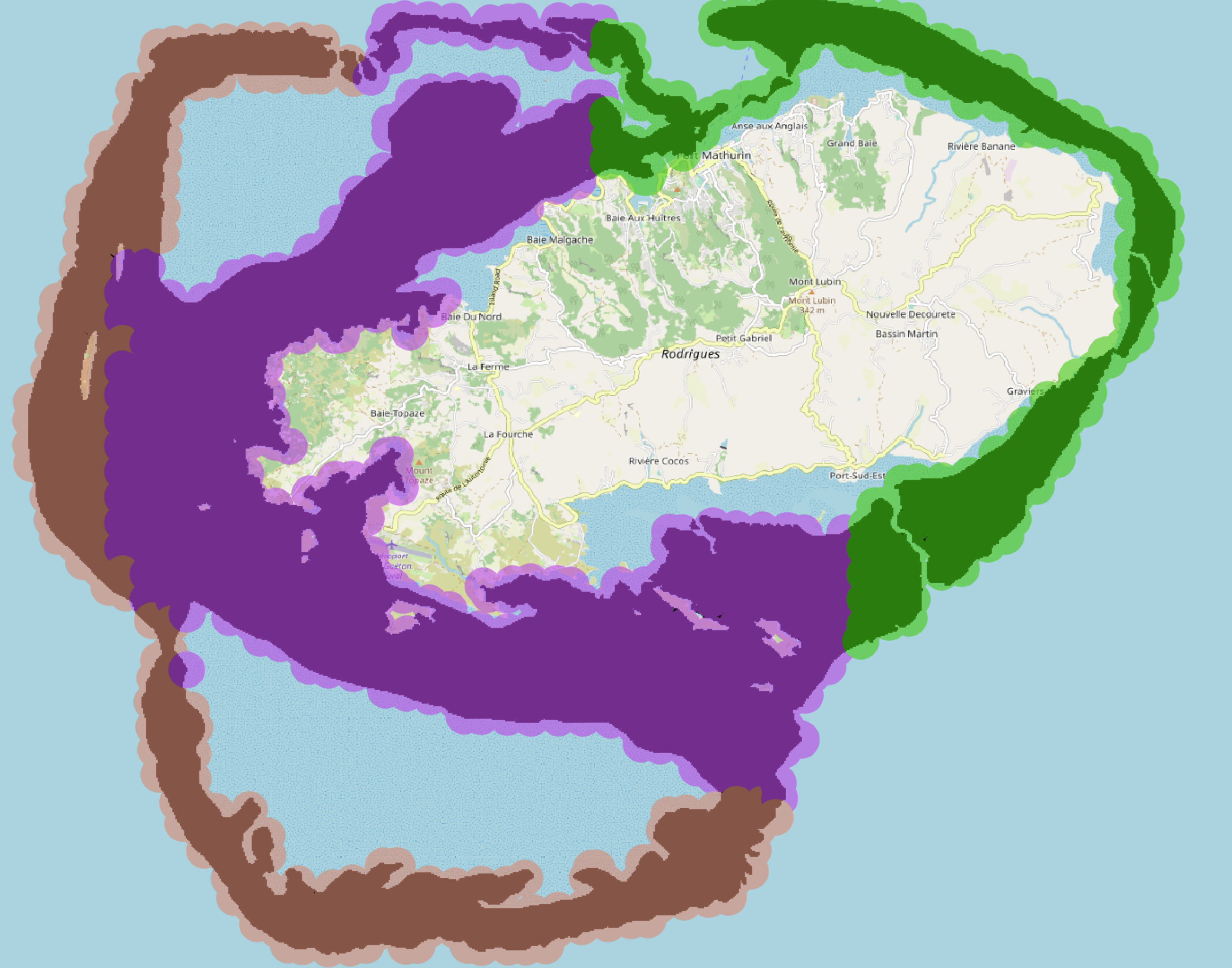

| ENV_zone | Degree heating week | SST st deviation | Chlorophyll mean | pH mean | Sea current velocity st. deviation | Depth in a 10 km buffer | Boat detection in a 10 km buffer | Population density in a 10 km buffer |
| --- | --- | --- | --- | --- | --- | --- | --- | --- |
| 1 | 0.727±0.0069 | 1.7±0.0026 | 0.227±0.053 | 8.07±5.2e-05 | 0.0555±0.015 | -197±150 | 0.000441±0.00034 | 53.9±24 |
| 2 | 0.73±0.009 | 1.7±0.0023 | 0.323±0.071 | 8.07±6e-05 | 0.0671±0.006 | -50.5±40 | 0.000901±0.00023 | 141±41 |
| 3 | 0.702±0.0057 | 1.69±0.0014 | 0.2±0.09 | 8.07±0 | 0.0607±0.0063 | -66.2±18 | 0.000709±0.00028 | 181±21 |

**Mahé, Seychelles**

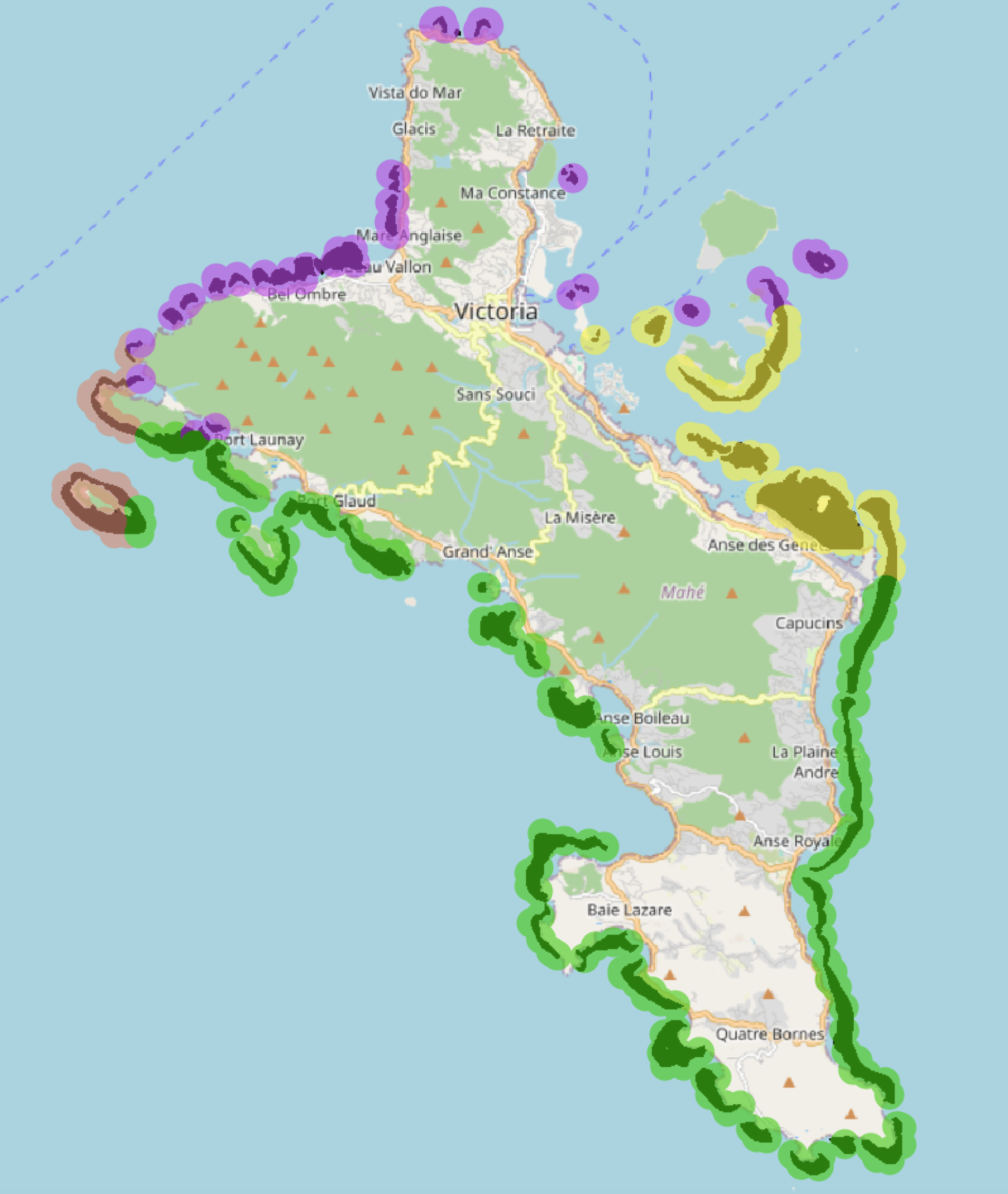

| ENV_zone | Degree heating week | SST st deviation | Chlorophyll mean | pH mean | Sea current velocity st. deviation | Depth in a 10 km buffer | Boat detection in a 10 km buffer | Population density in a 10 km buffer |
| --- | --- | --- | --- | --- | --- | --- | --- | --- |
| 1 | 0.268±0.0052 | 1.26±0.0047 | 0.225±0.0073 | 8.04±0 | 0.102±0.00058 | -44.7±1.1 | 5e-04±3e-04 | 142±35 |
| 2 | 0.276±0.0058 | 1.25±0.0035 | 0.296±0.03 | 8.04±5e-1 | 0.0993±0.018 | -45±6.4 | 0.0437±0.029 | 354±88 |
| 3 | 0.268±0.0045 | 1.27±0.0074 | 0.274±0.054 | 8.04±0 | 0.0948±0.013 | -39.3±5.9 | 0.00309±0.0024 | 228±90 |
| 4 | 0.274±0.0039 | 1.26±0.0048 | 0.371±0.077 | 8.04±0 | 0.0632±0 | -50±1.6 | 0.0587±0.007 | 431±59 |

**Praslin, Seychelles**

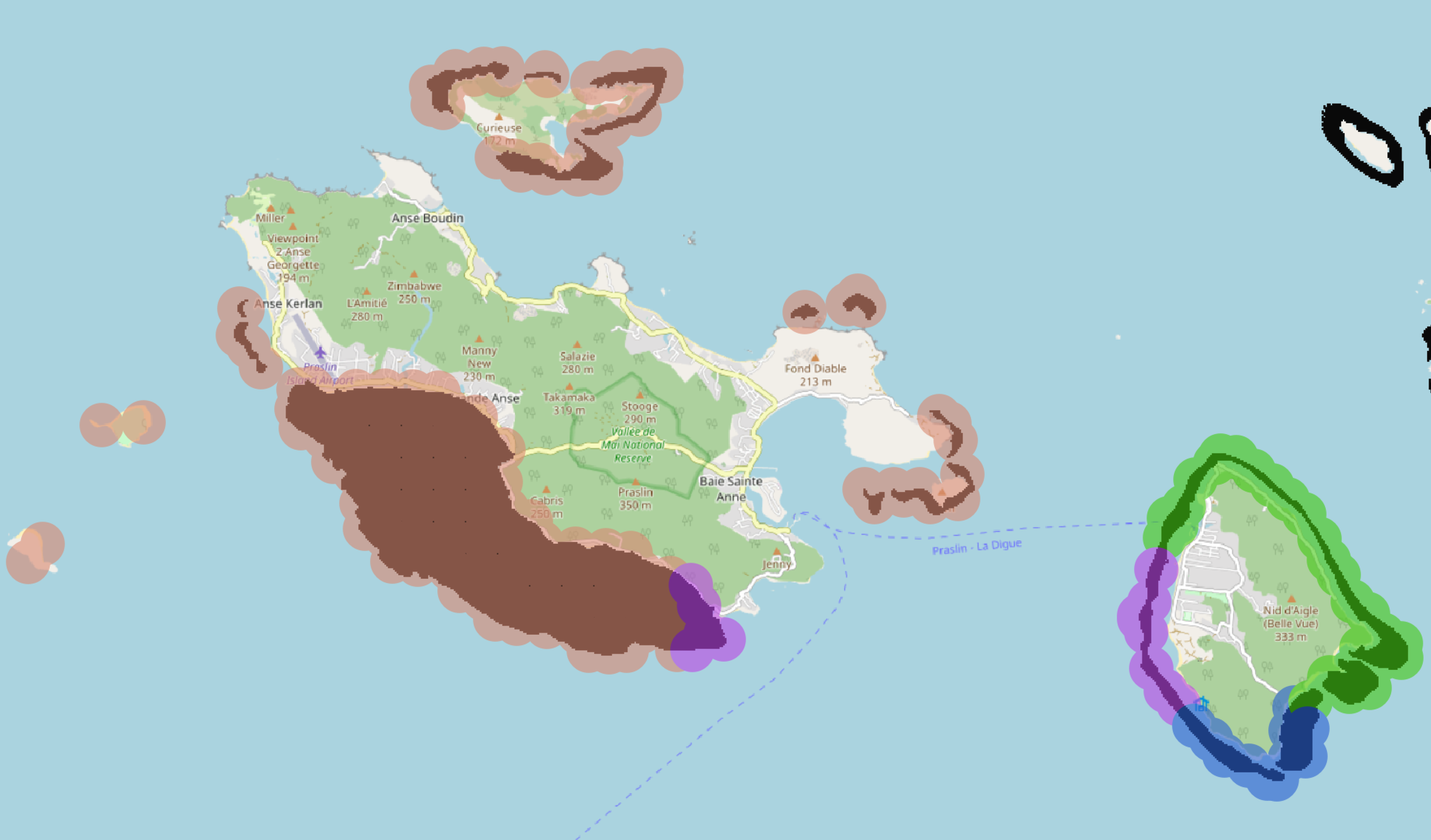

| ENV_zone | Degree heating week | SST st deviation | Chlorophyll mean | pH mean | Sea current velocity st. deviation | Depth in a 10 km buffer | Boat detection in a 10 km buffer | Population density in a 10 km buffer |
| --- | --- | --- | --- | --- | --- | --- | --- | --- |
| 1 | 0.26±0.0027 | 1.22±0.003 | 0.304±0.023 | 8.04±0 | 0.0731±0.0068 | -23.1±2.2 | 0.00602±0.0011 | 142±19 |
| 2 | 0.255±0.00063 | 1.22±0.0011 | 0.3±0.0026 | 8.04±0 | 0.0739±0.004 | -26±2.3 | 0.0052±0.0016 | 138±6.9 |
| 3 | 0.261±0.0018 | 1.21±0.0016 | 0.27±0.016 | 8.04±0 | 0.0888±0.0076 | -31±2.5 | 0.00317±0.00084 | 123±12 |
| 4 | 0.256±0 | 1.22±0 | 0.23±0.0035 | 8.04±0 | 0.115±0 | -30.8±0.92 | 0.00277±0.00046 | 123±2.5 |
